# Four phases and temporal threshold of population calcium response during locomotion in cortical astrocytes

**DOI:** 10.1101/2025.11.15.688660

**Authors:** Anna Fedotova, Alexey Brazhe, Alexey Semyanov

## Abstract

Calcium activity is a major form of astrocyte excitability essential to their function and plasticity. Astrocytic calcium events arise both spontaneously and in response to behaviour ^1^. In this study, we examined how these unitary events shape population-level calcium activity during quiescence and locomotion. Using two-photon imaging of astrocytic calcium in the primary somatosensory cortex (S1) of awake mice running on a freely rotating disk, we found that spontaneous fluctuations during quiescence were driven mainly by the co-occurrence of new calcium events rather than by their enlargement. During locomotion, population responses progressed through four phases: the emergence of new events, their merging into a superevent, fragmentation of this superevent, and an afterburst. New events appeared immediately at locomotion onset, indicating a rapid astrocytic response. The response also exhibited a temporal threshold of ∼5 seconds: when locomotion was shorter than this, the superevent phase failed to develop, and the population response was markedly reduced. In suprathreshold responses, the superevent displayed a ΔF/F intensity pattern that reproduced reliably across locomotion episodes. Hotspots within this pattern had shorter latencies and were also more active during quiescence, suggesting that the response is at least partly deterministic. Consistent with this, we were able to reproduce both the response phases and the superevent pattern using a dynamic mode decomposition with control (DMDc) model driven by synthetic or real locomotion inputs. Together, these findings dissect the population calcium response into quantifiable properties of unitary calcium events and reveal the formation and reproducibility of the superevent pattern, offering a potential framework for the astrocytic component of the engram ^2,3^.

---

Brain function emerges from the coordinated activity and interactions among diverse cell types - neurons, glial cells (astrocytes, microglia, and oligodendrocyte lineage cells), vascular cells, and non-cellular components such as the extracellular space and extracellular matrix. Together, these elements and interactions form the brain active milieu (BAM) ^4,5^. As a component of BAM, astrocytes play a significant role in intercellular communication, ion and neurotransmitter homeostasis, and metabolic processes ^6,7^. Astrocytes interact with virtually all other elements of the BAM - regulating microglial activation, modulating synaptic transmission and neuronal excitability, communicating with oligodendrocyte precursor cells and oligodendrocytes, and participating in neurovascular coupling. Astrocytes contribute to the remodelling of the extracellular space and release of extracellular matrix components ^8-10^.

Unlike neurons, astrocytes are electrically non-excitable cells. The principal form of their excitability and signalling is intracellular calcium activity ^1^. Astrocytic calcium events typically emerge at the periphery of the astrocytic domain and propagate to the soma, where they trigger larger somatic calcium surges ^11–16^. Integration of calcium events in astrocytes resembles synaptic integration in neurons, albeit with distinct spatiotemporal dynamics. While synaptic inputs in neurons are anatomically well-defined, the precise initiation sites of astrocytic calcium events remain a matter of debate. Some studies suggest that calcium signals consistently originate from specialised anatomical microdomains, while others report a more stochastic, spatially distributed activity pattern ^17–21^. In addition, astrocytic calcium events exhibit complex spatiotemporal dynamics, as they can expand, shrink, propagate, fragment, or merge, rendering the traditional region-of-interest (ROI)-based approach insufficient for their analysis ^17,18,22^.

Astrocytic calcium activity is closely linked to animal behaviour and arousal. During locomotion, astrocytic response in multiple brain regions is triggered through *locus coeruleus* projections, releasing noradrenaline and activating astrocytic α_1_-adrenergic receptors ^12,23–25^. Sensory inputs also contribute to astrocytic calcium responses. For example, combining visual stimuli with noradrenaline release or locomotion significantly increases astrocytic calcium activity in the visual cortex ^23,25^. These studies have suggested that arousal during locomotion amplifies astrocytic calcium response to sensory input.

In the hippocampus, astrocytic calcium transients occur during exploration and learning, and manipulating astrocyte calcium can affect memory formation and retrieval ^26^. A recent study has demonstrated that learning activates specific hippocampal astrocytic ensembles, characterised by elevated calcium signalling with greater amplitude and duration of calcium surges ^3^. Another study found that fear experience triggers astrocytic calcium and cAMP activity in the amygdala, driven by noradrenergic signalling ^2^. During fear memory recall, astrocytes showed increased frequency of population calcium responses and prolonged cAMP elevation. These findings suggest that astrocytic calcium, in addition to sensory input processing, can also actively shape cognitive function and contribute to the astrocytic component of BAM engram ^5^.

Although astrocytic calcium signalling is increasingly recognised as a key component of the BAM response to sensory and neuromodulatory input, the role of unitary calcium events in shaping astrocytic population activity remains poorly understood. In this study, we investigated how unitary calcium events contribute to astrocytic network activity in the somatosensory cortex of mice during quiescence and locomotion. During quiescence, population calcium fluctuations were primarily driven by the co-occurrence of new calcium events. The population response to locomotion progressed through four sequential phases: emergence of new calcium events, their merging into a superevent, fragmentation of the superevent, and an afterburst. Astrocytic response to locomotion also displayed a temporal threshold of 5 seconds. Shorter (subthreshold) episodes of locomotion induced only low-amplitude, spatially confined calcium responses that failed to develop the superevent phases. Superevents formed during suprathreshold locomotion episodes and persisted until the end of locomotion. It was characterised by a specific pattern of calcium signal intensities (ΔF/F), which may be instrumental for the astrocytic component of the BAM engram.

## Results

### Population calcium activity and unitary events in cortical astrocytes

We conducted two-photon calcium imaging in layer I astrocytes of the primary somatosensory cortex (S1) of head-fixed mice placed on a freely rotating disk (Figure 1a and Supplementary Video 1). The genetically encoded calcium sensor GCaMP6f was expressed in astrocytes following stereotaxic injection of the viral vector AAV9-gfaABC1D-GCaMP6f. After the injection, we attached a glass window and a metal head holder to the animal’s skull. After a recovery period of four to six weeks, the mice were handled for an additional week before the experiment.

**Figure 1:**
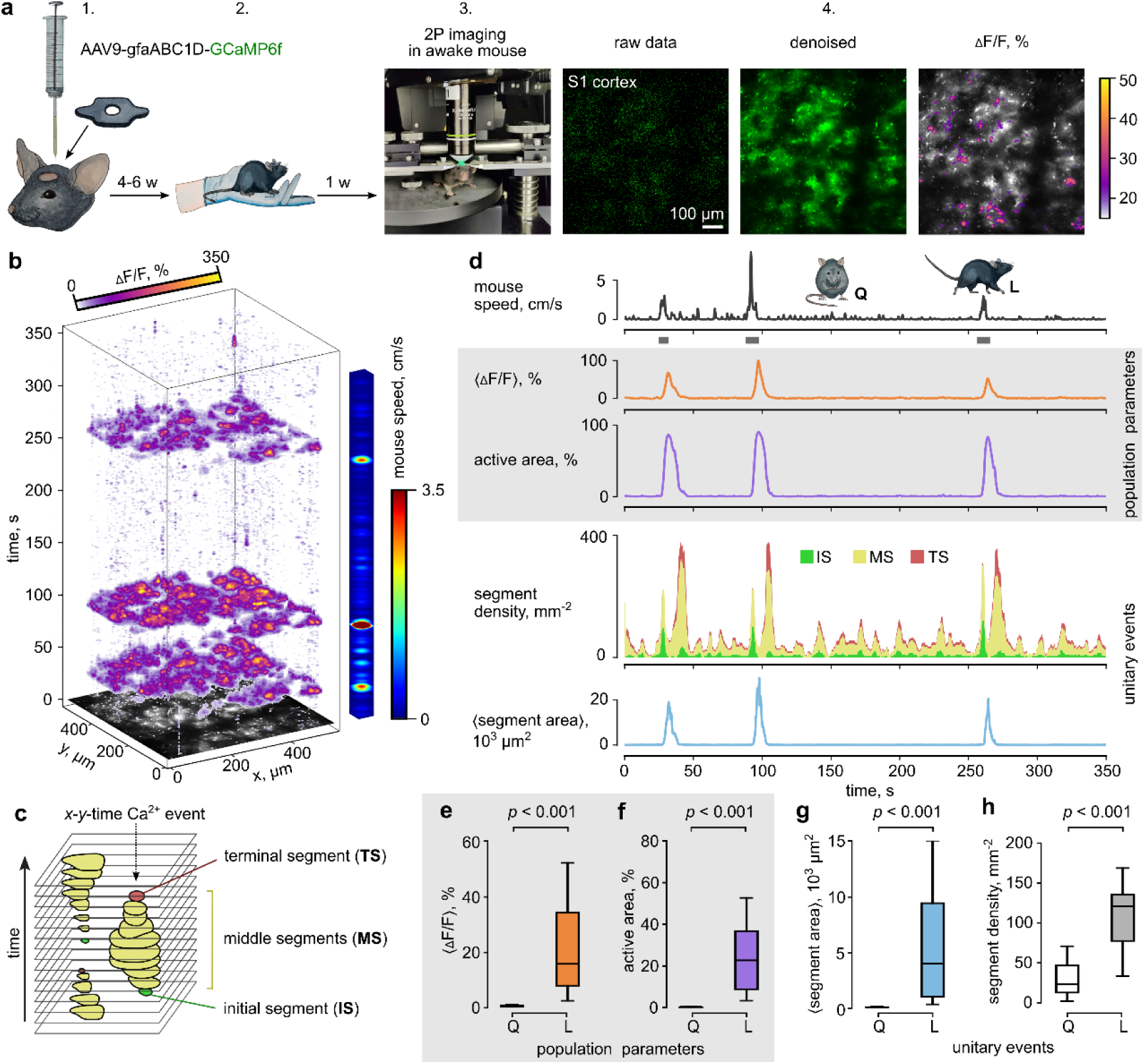
Population calcium activity and unitary events in cortical astrocytes. **a**. A schematic of the experimental procedure: 1. AAV injection for astrocyte-specific GCaMP6f expression in the somatosensory cortex (S1). 2. Habituation and handling of the mouse. 3. Calcium imaging in a head-fixed awake mouse on a freely rotating disk. 4. Image processing pipeline: from left to right - raw image, image denoising and motion correction, ΔF/F calculation for each pixel. **b.** A three-dimensional reconstruction (*x*-*y*-time) of ΔF/F that demonstrates the distribution of astrocytic calcium activity across the astrocytic network over time. The coloured bar on the right indicates the speed of the animal. **c.** Calcium events were reconstructed in space and time from individual segments overlapping in subsequent frames. Initial segments (IS, green), middle segments (MS, yellow), and terminal segments (TS, red) characterise the appearance, evolution, and termination of calcium events, respectively. **d.** Timecourses of mouse speed, population parameters (<ΔF/F> and active area, shown in grey), and unitary calcium events (segment density and <segment area>). The stacked area graph is colour-coded: green for IS, yellow for MS, and red for TS. Dark grey bars mark locomotion episodes (L), alternating with quiescence (Q). **e-h.** Summary data showing changes in population parameters (<ΔF/F> and active area, shown in grey), and unitary calcium events (segment density and <segment area>) averaged during quiescence (Q) and locomotion episodes (L) in individual imaging sessions. Data are presented as median [Q1; Q3] ± 1.5 IQR, with *p*-values obtained from a linear mixed model (LMM) based on 22 imaging sessions from 7 animals.

Recorded images were motion-corrected and denoised before the relative changes in fluorescence (ΔF/F) were calculated in each pixel (for further details, see the Methods section). Calcium events were defined both in space and time as adjacent pixels with suprathreshold ΔF/F values overlapping in subsequent frames ^18^. Consistent with previous studies, we observed scattered calcium events during animal quiescence, but large calcium responses during locomotion episodes (Figure 1b)^12,23–25^.

To characterise spatiotemporal dynamics of calcium events, their profiles in individual time frames were classified into three categories: initial segments (IS), which indicated the appearance of new events in a specific frame; terminal segments (TS), which marked the termination of events; and middle segments (MS), which characterised the spatiotemporal evolution of events (Figure 1c).

Calcium activity was characterised by two population parameters: the mean ΔF/F in the frame (<ΔF/F>) and the active area (Figure 1d) ^13^. Both parameters significantly increased during locomotion episodes: <ΔF/F> was 0.69 [0.55; 0.92] % during quiescence and 15.80 [7.99; 34.14] % during locomotion (n-numbers and p-values are present in corresponding figures; Figure 1e); the active area was 0.15 [0.08; 0.33] % of the fluorescent area during quiescence and 22.77 [8.96; 36.73] % during locomotion (Figure 1f).

The population calcium responses during locomotion episodes were characterised by an increase in the mean segment area (<segment area>), which was 0.06 [0.05; 0.08] × 10^3^ μm² during quiescence and 3.99 [1.02; 9.39] × 10^3^ μm² during locomotion (Figure 1d, g). Segment densities showed two notable peaks flanking the <segment area> peak (Figure 1d). The segment densities were significantly higher during locomotion episodes compared to quiescence, with values of 22.92 [12.28; 46.58] mm⁻² during quiescence and 120.15 [77.03; 134.82] mm⁻² during locomotion (Figure 1h). These results suggest that the population calcium response during locomotion is mediated by the appearance of new calcium events, which merge and then fragment again.

Next, we analysed in greater detail the contribution of individual calcium events to population calcium activity during both quiescence periods and locomotion episodes.

### Population calcium fluctuations during quiescence

The periods of animal quiescence were identified between locomotion episodes. During these periods, the animal made short movements on the disk that did not reach the threshold for classification as locomotion (see Method section, Figure 2a, b). While the animal was in a quiescent state, we also observed fluctuations in both <ΔF/F> and the active area of calcium activity, which showed little correlation with the short movements (correlation coefficient for <ΔF/F>: 0.05 [0.03; 0.08], and for active area: 0.12 [0.09; 0.17]; Extended Data Figure 1).

**Figure 2:**
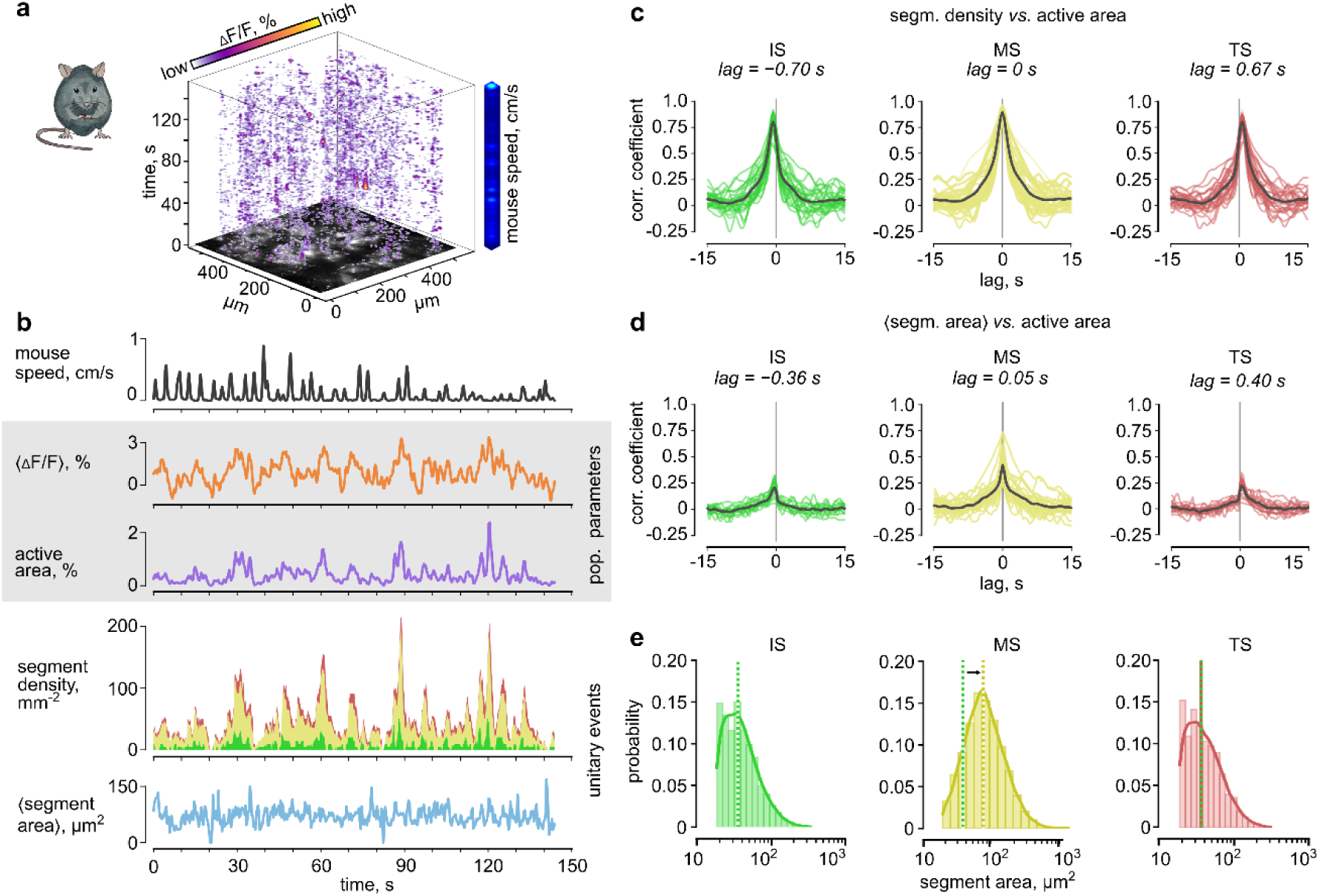
Population calcium fluctuations during mouse quiescence are driven by the co-occurrence of new calcium events. **a**. A three-dimensional reconstruction of astrocytic calcium activity during animal quiescence. The coloured bar on the right shows the speed of the animal. **b.** Timecourses of mouse speed, population parameters (<ΔF/F> and active area, shown in grey), and unitary calcium events (segment density and <segment area>) during mouse quiescence. The stacked area graph is colour-coded: green for IS, yellow for MS, and red for TS. **c.** Graphs of cross-correlation between segment density (IS, MS, and TS) and the active area. Vertical grey line marks zero lag. **d.** Graphs of cross-correlation between the <segment area> (IS, MS, and TS) and the active area. **e.** The distributions of segment areas (IS, MS, and TS) during quiescence for a single imaging session. Dotted lines indicate medians for IS (green), MS (yellow) and TS (red). Data in panels *c-e*: *n* = 26 periods of quiescence, 12 imaging sessions from 3 animals. For statistical comparison, see Extended Data Figure 2.

Fluctuations in the active area can originate from increases in the number of calcium events, their enlargement, or both. We detected a strong cross-correlation between segment density and active area (correlation coefficient for IS: 0.82 [0.75; 0.86], for MS: 0.90 [0.86; 0.93], and TS: 0.81 [0.76; 0.87]; Figure 2c; Extended Data Figure 2). In contrast, cross-correlation between <segment area> and active area was much weaker (correlation coefficient for IS: 0.20 [0.16; 0.27], for MS: 0.42 [0.33; 0.51], and TS: 0.23 [0.16; 0.28]; Figure 2d; Extended Data Figure 2). These results suggest that the appearance of new calcium events rather than their enlargement predominantly contributes to calcium activity fluctuations during quiescence.

However, the correlation coefficient for MS <segment area> was larger than that for IS and TS <segment area>, suggesting that event enlargement does play a role. Consistent with this observation, the distribution of MS areas was shifted to the right compared with those of IS and TS areas (IS areas: 35.8 [25.4; 53.1] μm², MS areas: 71.5 [47.3; 111.9] μm², TS areas: 35.8 [24.2; 55.4] μm²; Figure 2e).

### Four phases of the population calcium response during locomotion

Next, we analysed astrocytic calcium responses during locomotion episodes. Consistent with previous reports ^13,25^, the onset of population calcium response lagged the beginning of the locomotion episode and persisted for several seconds afterwards (Figure 3a, b). The time to peak (TTP) of <ΔF/F> from the beginning of locomotion was 6.30 [5.49; 8.32] s, and the TTP of the active area was 6.30 [5.49; 8.40] s (Figure 3c).

**Figure 3:**
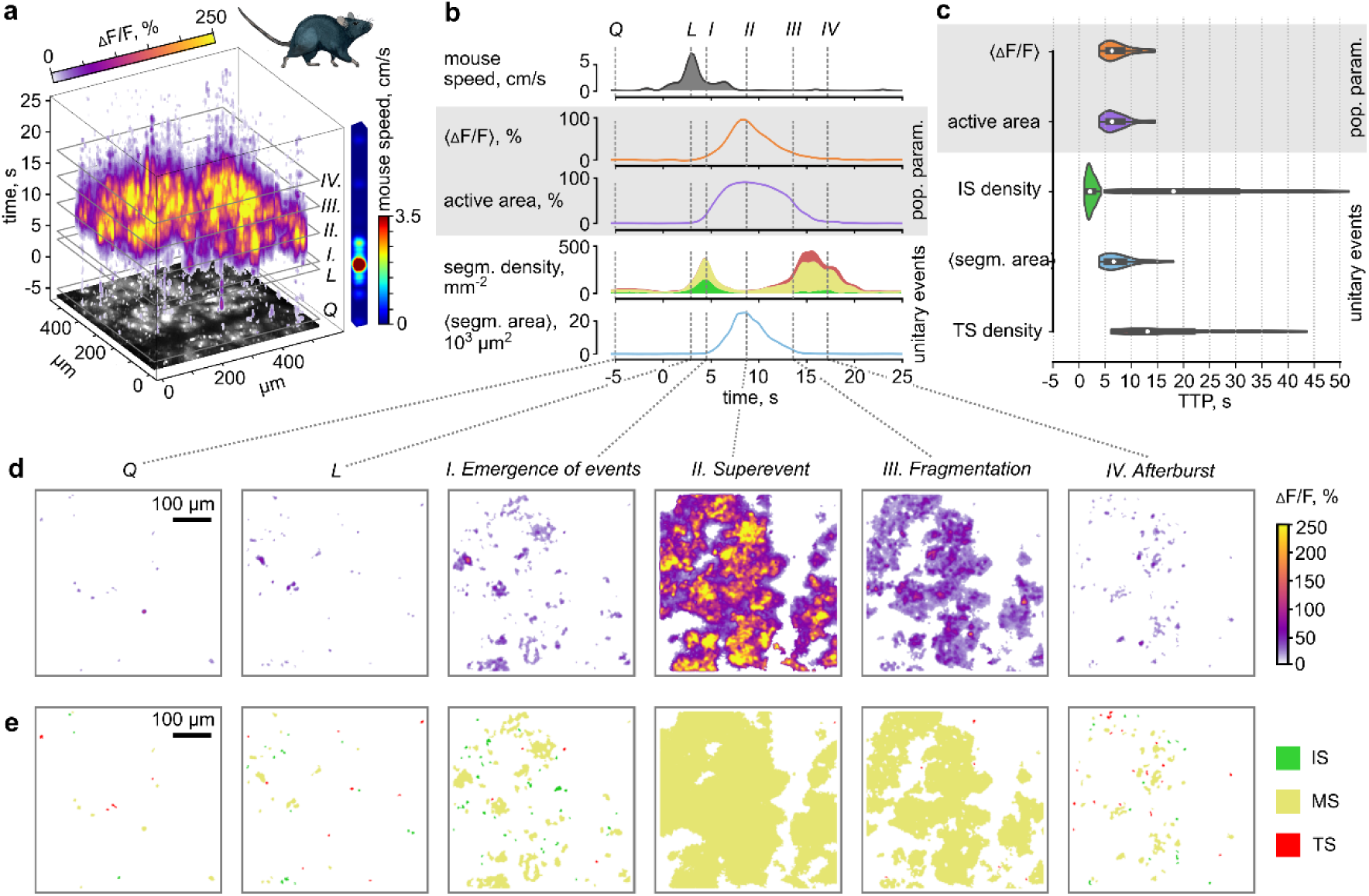
Four phases of population calcium response during locomotion. **a**. A three-dimensional reconstruction of astrocytic calcium activity during a locomotion episode. The coloured bar on the right shows the speed of the animal. **b.** Timecourses of mouse speed, population parameters (<ΔF/F> and active area, shown in grey), and unitary calcium events (segment density and <segment area>) during a locomotion episode. The stacked area graph is colour-coded: green for IS, yellow for MS, and red for TS. Dashed lines across the timecourses: Q for quiescence, L for peak speed, I, II, III, IV for the four phases of calcium response to locomotion. **c.** The distributions of TTPs for <ΔF/F>, active area, IS density, <segment area>, and TS density based on n = 38 episodes of locomotion in 13 imaging sessions from 7 animals. **d,e.** The frames show the spatial distribution of ΔF/F (*d*) and colour-coded segments (**e**): IS (green), MS (yellow), TS (red) at time points, marked with dashed lines in *b*.

In contrast to population parameters (<ΔF/F> and active area), the changes in unitary calcium events were much more complex. We identified four phases of such changes in the astrocytic response during a locomotion episode (Figure 3b-e, Supplementary Video 2).

The first phase began shortly after locomotion started and was characterised by the emergence of new calcium events indicated by a sharp peak in IS density. The TTP of IS density was 2.10 [1.62; 2.83] s. Thus, the astrocytic response to locomotion was much faster than that inferred from population parameters.

In the second phase, the events grew larger and began to merge, ultimately forming a superevent. The density of segments decreased, while the <segment area> increased. The TTP of <segment area> was 6.62 [5.25; 8.73] s. Around this time, the population parameter reached their maxima.

In the third phase, the superevent began to fragment into separate events that then terminated. The segment density increased again, and the TS density began to grow. These processes corresponded to the decay in population parameters.

In the fourth phase, we observed a peak in TS density, followed by a smaller peak in IS density. TTP of TS density was 13.09 [9.78; 21.81] s, and TTP of IS density was 18.10 [10.66; 30.38] s. This result suggests that, despite the termination of previous calcium events, new calcium events appeared. This dynamic, attributed to residual astrocytic activity, was termed an afterburst.

### The temporal threshold for the population calcium response

The type, duration, and intensity of the stimulus strongly influence astrocytic calcium responses ^11^. It has been demonstrated that noradrenaline signalling in astrocytes integrates over time ^27^. The authors showed that brief optogenetic stimulation of noradrenergic fibres (lasting less than 5 seconds) triggers a minimal population calcium response in astrocytes, whereas longer stimulations elicit significantly larger responses. Here, we examined how population calcium responses depend on locomotion duration, because locomotion is naturally accompanied by activation of noradrenergic projections, leading to endogenous release of noradrenaline ^25,28^.

We observed that short episodes of locomotion trigger subthreshold population responses. These responses were characterised by a small <ΔF/F> and by a small active area (Figure 4a). Our analysis revealed two distinct clusters in the scatter plot of peak active area versus peak <ΔF/F> (Figure 4b), corresponding to subthreshold and suprathreshold responses. Based on these two clusters, we defined the threshold at a peak active area of 50% and a peak <ΔF/F> at 25%.

**Figure 4:**
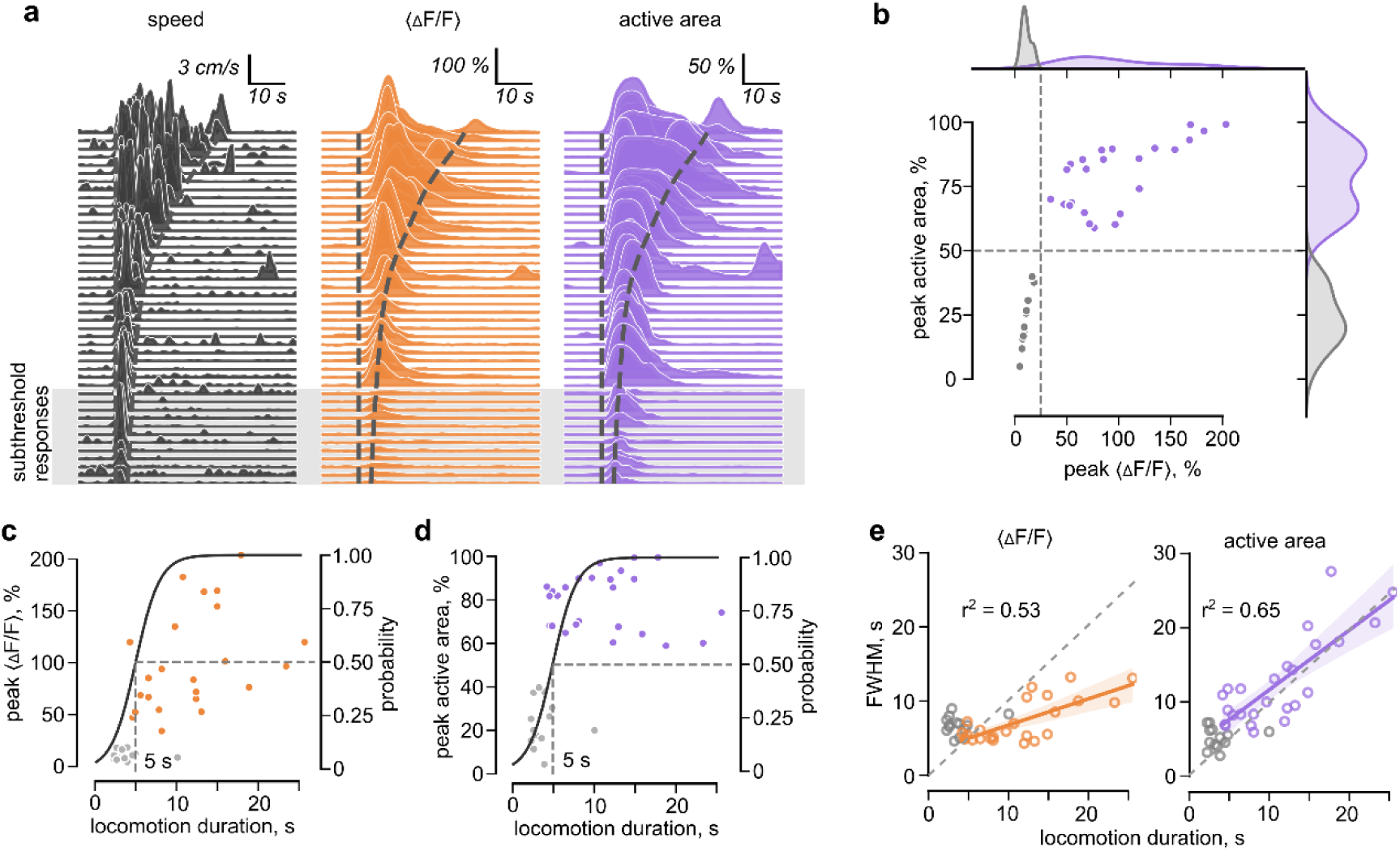
The temporal threshold for the population calcium response. **a.** The waterfall plots display the timecourses of animal speed (black), <ΔF/F> (orange), and active area (violet), sorted by the duration of locomotion episodes. Each curve represents a single locomotion episode. Dashed lines indicate the beginning and end of the episode. Subthreshold responses are shown in grey. **b.** The relationship between the peak active area and the peak <ΔF/F>. The responses were considered suprathreshold when the peak active area exceeded 50% and the peak <ΔF/F> exceeded 25%, as indicated by the dashed lines. Grey circles and marginal distributions represent subthreshold responses, while violet circles and distributions represent suprathreshold responses. **c.** The relationship between peak <ΔF/F> and the locomotion duration, overlaid with a binary logistic regression model of suprathreshold response probability (black curve, *right axis*). Grey circles represent subthreshold responses, coloured circles represent suprathreshold responses. The dashed lines indicate a 50% probability of a suprathreshold response, along with the corresponding locomotion duration of 5 s. **d.** The same as panel ***c***, but for the peak active area. **e.** The relationships between the full width at half maximum (FWHM) of <ΔF/F> and locomotion duration (*left graph*), and the FWHM of active area and locomotion duration (*right graph*). Grey circles represent subthreshold responses, coloured circles represent suprathreshold responses. r^2^ is the coefficient of determination. The dashed lines indicate FWHM equal to the locomotion duration. Data: n = 38 episodes of locomotion in 13 imaging sessions from 7 animals

To determine the temporal threshold, we plotted the distributions of subthreshold and suprathreshold responses against the duration of locomotion, using peak <ΔF/F> (Figure 4c) and peak active area (Figure 4d) as metrics. We defined the temporal threshold as the locomotion duration necessary to generate suprathreshold responses with a probability of 0.5. Similar to the optogenetic stimulation of noradrenergic fibres, we identified the temporal threshold at 5 seconds ^27^.

Interestingly, suprathreshold locomotion episodes displayed distinctive dynamics of <ΔF/F> and active area (Figure 4a, e). The <ΔF/F> began to peak soon after locomotion onset and subsequently decreased. The active area reached its peak around the same time (Figure 3c) and remained at a high level until the locomotion finished. Consequently, the full width at half maximum (FWHM) of <ΔF/F> did not increase as much with locomotion duration as the FWHM of the active area. These observations suggest that while the calcium response remains spatially preserved throughout the locomotion episode, its amplitude declines over time. This decline may be attributed to the depletion of endogenous calcium stores or the desensitisation of α1-adrenoreceptors ^13,29^.

### The temporal threshold is linked to the superevent phase

To understand the mechanism behind the temporal threshold, we analysed the phases of population responses during locomotion episodes of varying durations (Figure 5a). In the first phase, new calcium events emerged in both subthreshold and suprathreshold responses. The peak of IS density consistently occurred at the beginning of each locomotion episode, and the TTP of IS density did not significantly vary with the duration of locomotion (Figure 5b). However, the peak of IS density was significantly larger in suprathreshold responses (118.3 [110.5, 144,9] mm^-2^) compared to subthreshold responses (78.2 [64.9, 85.5] mm^-2^; Figure 5c, d).

**Figure 5:**
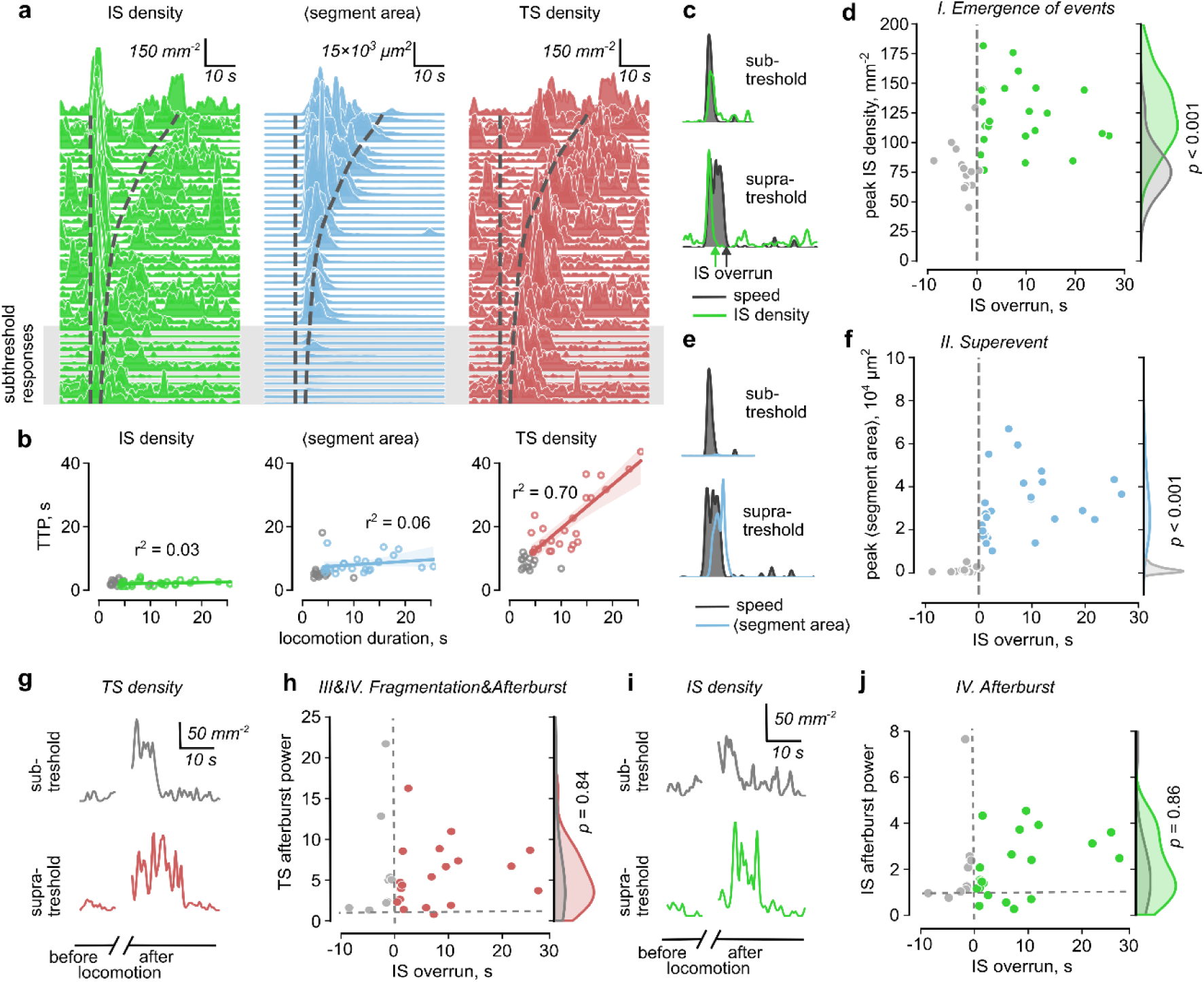
Dynamics of calcium events during subthreshold and suprathreshold responses. **a.** The waterfall plots display the timecourses of IS density (green), <segment area> (blue), and TS density (red), sorted by the duration of locomotion episodes. Each curve represents a single locomotion episode. Dashed lines indicate the beginning and end of the episode. Subthreshold responses are shown in grey. **b.** The relationships between the TTP for IS density (*left*), TTP for <segment area> (*middle*), and TTP for TS density (*right*) and locomotion duration. Grey circles represent subthreshold responses, coloured circles represent suprathreshold responses. r^2^ is the coefficient of determination. **c.** Sample traces of IS density (green) and animal speed (black with shaded area) for subthreshold (*top*) and suprathreshold (*bottom*) responses. The green arrow marks the end of the IS density peak, and the black arrow marks the end of the locomotion episode. The interval between these arrows represents the IS overrun. **d.** The relationship between peak IS density (phase I, emergence of events) and IS overrun. Negative IS overruns indicate that the animal stopped running before the IS density peak finished. The dashed line indicates zero IS overrun separating subthreshold and suprathreshold responses. Grey circles and grey marginal distribution represent subthreshold responses, green circles and green distribution represent suprathreshold responses. **e.** Sample traces of <segment area> (blue) and animal speed (black with shaded area) for subthreshold (*top*) and suprathreshold (*bottom*) responses. **f.** The relationship between peak <segment area> (phase II, superevent) and IS overrun. The dashed line indicates zero IS overrun. Grey circles and grey marginal distribution represent subthreshold responses, blue circles and blue distribution represent suprathreshold responses. **g.** Sample traces of TS density before and after the locomotion episode for subthreshold (grey, *top*) and suprathreshold (green, *bottom*) responses. **h.** The relationship between TS afterburst power (phases III/IV, fragmentation and afterburst) and IS overrun. The vertical dashed line indicates zero IS overrun, the horizontal dashed line indicates TS afterburst power = 1. Grey circles and grey marginal distribution represent subthreshold responses, red circles and red distribution represent suprathreshold responses. **i.** Sample traces of IS density before and after the locomotion episode for subthreshold (grey, *top*) and suprathreshold (green, *bottom*) responses. **j.** The relationship between IS afterburst power (phase IV, afterburst) and IS overrun. The vertical dashed line indicates zero IS overrun, the horizontal dashed line indicates TS afterburst power = 1. Grey circles and grey marginal distribution represent subthreshold responses, red circles and red distribution represent suprathreshold responses. Data: *p*-values are LMM; 9 subthreshold and 21 suprathreshold responses in 12 imaging sessions from 7 animals

We also noticed that suprathreshold responses only developed when the locomotion episode continued after the end of the IS density peak (emergence of events phase I). We introduced a parameter called ‘IS overrun’ to quantify the interval between the end of the IS density peak and the end of animal locomotion. Negative IS overruns corresponded to subthreshold responses, positive to subthreshold ones (Figure 5d).

In suprathreshold responses, the peak of IS density was followed by a peak of <segment area> (superevent phase II). The TTP of the <segment area> also did not show a clear dependence on locomotion duration (Figure 5b). However, the <segment area> peak was strongly suppressed in subthreshold responses, suggesting that the superevent did not develop. The <segment area> peak was 0.07 [0.05, 0.22] × 10^4^ µm^2^ in subthreshold responses and 3.17 [2.00, 3.74] × 10^4^ µm^2^ in suprathreshold responses (Figures 5e, f). Thus, the emergence of new calcium events during locomotion does not lead to the formation of a superevent if the animal stops running before the end of the IS density peak. Consequently, the population parameters, <ΔF/F> and active area, also get suppressed.

Despite the differences in the superevent phase II, the fragmentation phase III and the afterburst phase IV were clearly pronounced in both subthreshold and suprathreshold responses. At the end of locomotion episodes, calcium events started to fragment and terminate. Thus, the TTP of TS density exceeded the duration of locomotion (Figure 5b).

To analyse event termination during phases III and IV, we calculated a ratio between the mean TS density after locomotion and the mean TS density before locomotion, which we termed TS afterburst power. The TS afterburst power was not significantly different between subthreshold (4.98 [2.19, 5.32]) and suprathreshold (4.37[2.58, 7.35]) responses (Figures 5g, h).

To examine the emergence of new events during the afterburst phase IV, we calculated a ratio between the mean IS density after locomotion and the mean IS density before locomotion, which we termed IS afterburst power. The IS afterburst power was not significantly different between subthreshold (1.26 [1.03, 2.38]) and suprathreshold (1.54 [1.16, 3.12]) responses (Figures 5i, j).

In summary, we show that the temporal threshold of the population calcium response is reached only after the animal surpasses the emergence of events phase I. Continued locomotion is required for these events to merge into a superevent. The fragmentation phase III and afterburst phase IV begin once locomotion ends and do not differ substantially between subthreshold and suprathreshold responses. Their characteristics are likely regulated by factors other than locomotion duration.

### Spatiotemporal pattern of population response during locomotion and its reproducibility

Although our primary goal was to characterise population calcium activity in astrocytes through changes in unitary calcium events, the superevent phase II represents a point of singularity. At this phase, individual events merge, and the system can no longer be described in terms of segment density or <segment area>. Nonetheless, the ΔF/F profile during the superevent phase II appeared spatially inhomogeneous.

To characterise the ΔF/F profile, we plotted calcium activity timecourses in each pixel (Figure 6a,b). Indeed, the maximal values of ΔF/F (ΔF/F_max_) of these timecourses were achieved during the superevent phase II (Figure 6b). Thus, the spatial distribution of ΔF/F_max_ can be used to characterise the ΔF/F profile during this phase.

**Figure 6:**
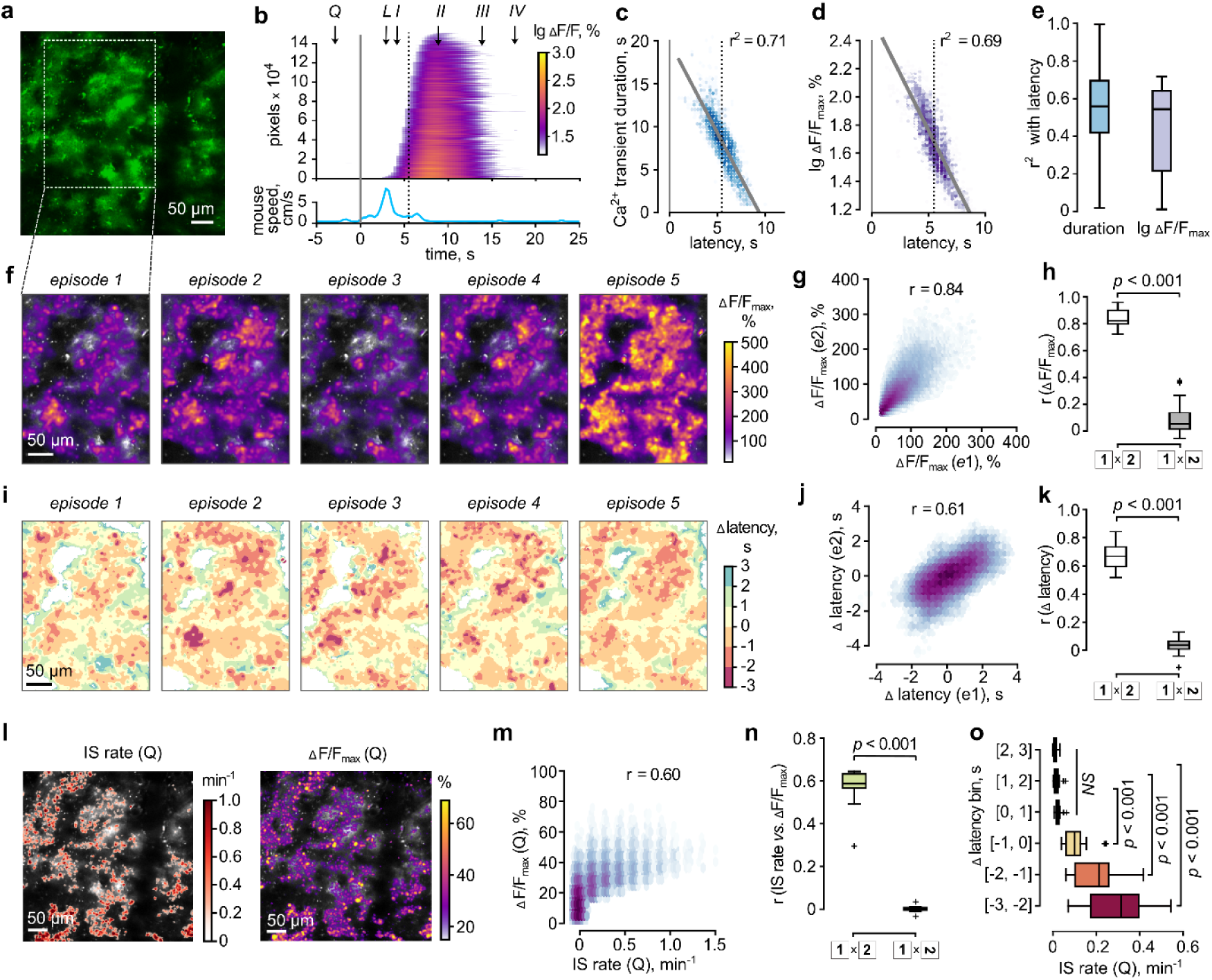
The pattern of population calcium responses during locomotion is reproducible in space in time in space and time. **a.** The sample frame illustrates a cortical area where calcium imaging was performed. The green regions indicate the expression of GCaMP6 in astrocytes. The boxed area highlights a region shown in panels *f* and *i*. **b.** The top of the panel displays calcium transients in each active pixel, sorted by their latency from the onset of locomotion (0 s), while the bottom shows the locomotion episode. The grey line marks the onset of locomotion; the dotted line indicates the median latency of the calcium transients; the arrows indicate phases of population response identical to Figure 3b. **c,d.** The relationships between duration and latency (*c*) and between lg ΔF/Fmax and latency (*d*) of calcium transients in individual pixels of one locomotion episode. The ’r^2^’ is the coefficient of determination. The vertical lines are the same as in panel *b*. **e.** The summary of r^2^ with latency for duration and lg ΔF/Fmax across multiple locomotion episodes. **f.** Maps of ΔF/Fmax of calcium transients in individual pixels in five sequential episodes of locomotion. **g.** The relationship between ΔF/Fmax in corresponding pixels in two episodes of locomotion, e1 and e2 (episodes one and two). ‘r’ is Pearson’s correlation coefficient. **h.** The summary of the r(ΔF/Fmax) in 24 pairs of locomotion episodes for co-oriented maps and when one map was rotated by 90 degrees. **i.** Maps of Δ latencies are shown for the same episodes of locomotion as in panel *f*. **j.** The relationship between Δ latencies in corresponding pixels in two episodes of locomotion. **k.** The summary of r(Δ latencies) in 24 pairs of locomotion episodes for co-oriented maps and when one map was rotated by 90 degrees. **l.** Maps of the IS rate (*left*) and ΔF/Fmax (*right*) in quiescence (Q). **m.** The relationship between ΔF/Fmax and IS rate in corresponding pixels based on the maps presented in panel *l*. **n.** The summary of r(IS rate vs. ΔF/Fmax) in 9 pairs for co-oriented maps and when one map was rotated by 90 degrees. **o.** The relationship between Δ latency during locomotion (binned in 1-second intervals) and IS rate during quiescence. Data are presented as median [Q1; Q3] ± 1.5 IQR, *p*-values are LMM.

The onset of calcium transients varied across individual pixels, with a median latency of 3.39 [2.91; 3.88] s from the start of locomotion (n = 44 locomotion episodes). Transients with shorter latencies exhibited longer durations and higher ΔF/Fₘₐₓ values. Specifically, transient duration was inversely correlated with latency (r² = 0.55 [0.41; 0.69]; Figure 6c, e), as was log-transformed ΔF/Fₘₐₓ (r² = 0.54 [0.21; 0.64]; Figure 6d, e). These findings indicate that during a locomotion episode, calcium events initiate in hotspots where calcium signals are larger and longer. Subsequently, calcium activity propagates to adjacent regions of the astrocytic network.

To test whether the hotspots are reproduced across different locomotion episodes, we mapped ΔF/F_max_ for each episode of locomotion (Figure 6f). We compared ΔF/F_max_ in corresponding pixels between pairs of maps from the same imaging session (Figure 6g and Extended Data Figure 3). Pearson’s correlation coefficient was 0.82 [0.80; 0.90] when the maps were co-oriented, and it dropped to 0.05 [0.02; 0.14] when one map was rotated by 90 degrees (Figure 6h).

Similarly, we mapped the Δ latencies (the latencies of calcium transients in individual pixels relative to the median latency of the response) during each locomotion episode (Figure 6i). We compared the Δ latencies between pairs of maps from the same imaging session (Figure 6j and Extended Data Figure 4). Pearson’s correlation coefficient was 0.67 [0.60; 0.74] for co-oriented maps and 0.04 [0.01; 0.07] when one map was rotated by 90 degrees (Figure 6k). These results indicate that the spatiotemporal profile of population calcium responses in cortical astrocytes is highly reproducible across locomotion episodes.

The reproducible profile may reflect the distribution of local synapses or noradrenergic projections, as well as the specific properties and morphological organisation of the astrocytes themselves. In either case, one could expect that calcium events will also emerge more frequently in the same hotspots during quiescent period calcium activity fluctuations.

To test this possibility, we mapped the IS rate (the number of IS per second) and ΔF/F_max_ during quiescence and found a significant correlation between the two maps (Figure 6l, m). Pearson’s correlation coefficient between the IS rate and ΔF/F_max_ was 0.58 [0.57, 0.63] for co-oriented maps and 0.004 [-0.004, 0.008] when one map was rotated by 90 degrees (Figure 6n). Thus, like during locomotion, calcium activity was more frequent and larger in specific hotspots during quiescent periods.

Finally, we asked if the hotspots active during quiescent periods are the same hotspots involved in population response during locomotion. We found that the pixels with higher IS rates during quiescence also had shorter latencies in population responses during locomotion (Figure 6o). This finding suggests that activity hotspots observed during quiescent periods indeed coincide with those seen during locomotion. By analysing the frequency of calcium activity in individual pixels during quiescence, one can predict the spatiotemporal properties of calcium response during locomotion, pointing to its deterministic nature.

### Dynamic mode decomposition model with control

Our experimental results suggest that the astrocytic calcium response during locomotion could represent a low-dimensional deterministic dynamical system. To test this hypothesis, we applied dynamic mode decomposition (DMD) ^30^, which fits linear dynamical systems to time-series data. We assumed that calcium dynamics is a stable system driven by locomotion and thus used the DMD with control (DMDc) variant ^31^. DMDc searches for a pair of linear operators *A* and *B* that predict the state of the system in the next time moment *t+δt* [F(*t+δt*)] given its current state at time *t* [F(*t*)] and the state of a control parameter [C(*t*)]:

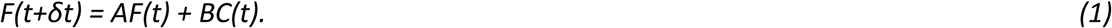

Matrix operator *A* is solved in the form of a combination of a low number of eigenmodes. Thus, starting from an initial state *F(t = 0),* one can simulate the F(*t*) trajectory in response to the recorded or synthesised realisation of control parameter C(*t*) by iteratively applying the estimates of operators A and B obtained from the fitting of prior experimental data. In the imaging data, we treated ΔF/F frames as system state F(*t*) and pairs of instantaneous mouse speed and acceleration values as control parameter C(*t*) (Extended Data Figure 5).

DMDc simulations using a real locomotion profile (shown in Figure 3b) for C(*t*) yielded a calcium response with similar spatiotemporal characteristics to the real astrocytic population response during this locomotion episode (compare Figure 7a to Figure 3a). DMDc model demonstrated similar spatiotemporal characteristics of the astrocytic population response to a locomotion episode, such as the shape and the delayed onset of both <ΔF/F> and active area changes.

**Figure 7:**
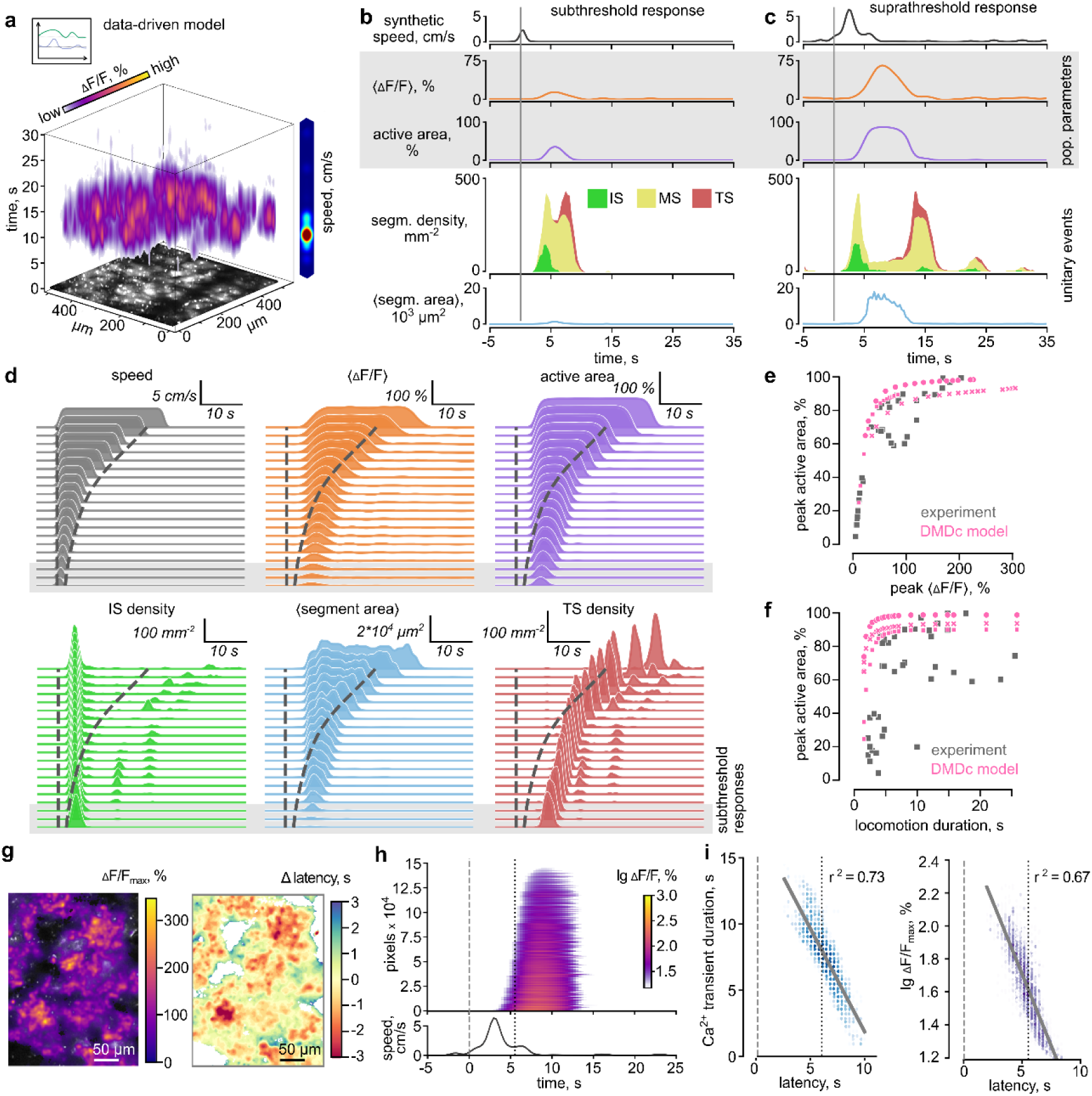
The data-driven DMDc model reproduces key features of population response in astrocytes. **a**. A three-dimensional reconstruction of astrocytic calcium activity during a locomotion episode based on the DMDc model. The coloured bar on the right shows the speed of the animal. **b,c.** Timecourses of mouse speed, population parameters (<ΔF/F> and active area, shown in grey), and unitary calcium events (segment density and <segment area>) during locomotion episode for subthreshold (**b**) and suprathreshold (**c**) responses based on DMDc model. Stacked area graph colour coding: green for IS, yellow for MS, red for TS. The vertical grey line indicates the onset of locomotion. **d.** The waterfall plots display the simulated timecourses of animal speed (grey), <ΔF/F> (orange), active area (violet), IS density (green), <segment area> (blue), and TS density (red), sorted by the duration of locomotion episodes. Each curve represents a single locomotion episode. Dashed lines indicate the beginning and end of the episode. Subthreshold responses are shown in the greyed area. **e.** The relationships between the peak active area and the peak <ΔF/F> for experimental data (grey squares, the data identical to Figure 4b) and generated with three runs of the DMDc model (pink circles, pink squares, and pink crosses). **f.** The relationships between the peak active area and locomotion duration for experimental data (grey squares, the data identical to Figure 4d) and generated with three runs of the DMDc model (pink circles, pink squares, and pink crosses). **g.** Maps of ΔF/F_max_ (*left*) and Δ latencies (*right*) of calcium transients in individual pixels in one synthetic episode of locomotion. **h.** The top of the panel displays calcium transients in each active pixel, sorted by their latency from the onset of locomotion; the bottom shows the locomotion episode used for the DMDc model. The dashed line marks the onset of locomotion, and the dotted line indicates the median latency of the calcium transients. **i.** The relationships between duration and latency (*left*) and between lg ΔF/Fmax and latency (*right*) of calcium transients in individual pixels. The ’r^2^’ represents the coefficient of determination, and vertical lines have the same meaning as indicated in panel *h*.

Strikingly, DMDc reproduced not only the four phases of population response but also subthreshold and suprathreshold population responses depending on locomotion duration (Figure 7b, c). Both subthreshold and suprathreshold responses exhibited phase I (emergence of events) characterised by a pronounced IS density peak. Then, in suprathreshold response, the events merged to form a superevent phase II, characterised by a substantial decrease in segment density with a large increase in segment area. The superevent phase II was not pronounced in the subthreshold response. The fragmentation phase III was strongly expressed in both types of responses. In contrast to experimental data, the afterburst phase IV was observed only in suprathreshold responses and characterised by several peaks of IS and TS densities.

The lack of multiple peaks in experimental afterburst may be explained by the refractoriness of calcium responses in astrocytic networks ^13,32^. The oscillatory modes revealed by the DMDc model may also be characteristic of a real astrocytic network, but refractoriness does not permit such oscillations.

The refractoriness of calcium responses is likely to have a complex nature, involving the depletion of calcium stores as well as the inactivation of receptors and signalling pathways. The inability of the astrocytic network to maintain a high level of calcium response for an extended period has been previously reported ^16^ and is evident in our experiments, as manifested by the decline of <ΔF/F> during prolonged episodes of locomotion (Figure 4a). The DMDc model did not demonstrate such a decline in <ΔF/F> (Figure 7d). We synthesised episodes of locomotion of different durations and observed sustained increases in <ΔF/F> and active area for the entire periods of locomotion. This result suggests that the DMDc model fails to reproduce the decline and refractoriness of astrocytic calcium responses to locomotion.

Set aside the lack of decline in <ΔF/F>, the relationships between peak <ΔF/F> and peak active area obtained with the DMDc model closely resembled the relationship for experimental data (Figure 7e). Similarly, subthreshold events generated with the DMDc model for short episodes of locomotion, as in experimental data (Figure 7f)

The parameters of unitary calcium events generated with the DMDc model quite closely reproduced the parameters in the real calcium response during locomotion episodes of different duration (Figure 7d). The peak of IS density during phase I was always generated at the beginning of locomotion. It was followed by the peak of <segment density> (superevent phase II). The peak of TS density is generated after locomotion ended (phases III and IV).

Finally, we analysed the spatiotemporal properties of the population responses generated by the DMDc model. The DMDc model produced spatial distributions of ΔF/F_max_ and Δ latencies similar to those observed in experimental data (compare Figure 7g to Figure 6f,i; Extended Data Figure 6). Like in experimental data, pixels with shorter latencies exhibited longer durations of calcium transients and greater amplitudes (Figure 7h,i).

These results suggest that the DMDc model essentially reproduces experimentally observed astrocytic calcium responses to locomotion episodes. Thus, the astrocytic network can be considered as a deterministic system where we can predict the response based on the specific input. However, DMDc does not capture the decay of calcium response that affects calcium dynamics at prolonged locomotion, which may play a role in the physiological outcome of astrocytic response.

## Discussion

We investigated how unitary calcium events contribute to population calcium activity in cortical astrocytes. During animal quiescence, we observed fluctuations of calcium activity due to the co-appearance of new calcium events. What causes such a simultaneous appearance of calcium events scattered throughout the astrocytic network remains unclear. Astrocyte-to-astrocyte signalling is not fast enough to trigger synchronised calcium elevation in distal parts. However, calcium fluctuations may be a response to slow fluctuations of adrenergic fibre activity observed during animal quiescence ^27,28,33^. Astrocytes can also respond to activity of local neuronal networks in the somatosensory cortex, which remain active during animal quiescence ^13,34,35^. Finally, astrocytes can respond to mechanical stimulation by parenchymal blood vessels through a transient receptor potential vanilloid subtype 4 (TRPV4) ^36,37^ or the mechano-gated Piezo1 channel ^38^.

Apart from the mechanism, another important question is the physiological role of population calcium fluctuations in astrocytes. It has been shown that spontaneous calcium events in the perirhinal cortex modulate synaptic strengthening and serve as a stochastic functional signal for memory consolidation ^39^. Spontaneous calcium activity in astrocytes may also play a role in vasomotion ^37^, hence regulating brain perfusion and glymphatic clearance. However, a recent report suggests that astrocytic cAMP rather than calcium is involved in the regulation of cerebellar blood flow ^40^. Astrocytic calcium has been implicated in the release of gliotransmitters ^41^. Through gliotransmission, calcium fluctuations can synchronise astrocytic networks, facilitating communication across large brain regions and coordinating neuronal network activity during tasks like sleep and attention ^42^. Calcium also regulates astrocytic morphology, which in turn affects synaptic coverage, glutamate uptake and ionic homeostasis ^43,44^. Thus, calcium fluctuations may maintain the morphological complexity of astrocytes and homeostatic support of the brain tissue. Finally, calcium regulates metabolic processes in astrocytes ^45,46^. That, in turn, may support the brain energy metabolism.

Following analysis of calcium fluctuations during animal quiescence, we investigated the population calcium response during locomotion. At first, this response was characterised by population parameters, such as <ΔF/F> and active area. Both parameters increased with a delay of a few seconds after the onset of locomotion, supporting the notion that astrocytes are slow responders as compared to neurons ^13,47–49^. Neurons respond rapidly during locomotion and are involved in the real-time processing of sensory and intrinsic (e.g., motor command, memories) brain inputs. The delayed astrocytic response contributes to slower processes, such as synaptic plasticity, metabolic processes, and restoration of ionic gradients.

On the other hand, multiple reports suggest that local calcium responses may appear on the fast time scale, and astrocytes can participate in sensory information processing ^50–52^. Such local responses may be undetectable in the population parameters but pronounced in changes of unitary calcium events during locomotion. Indeed, we observed an increase in IS density immediately after the beginning of locomotion (Figure 3c). The emergence of new calcium events marked phase I of the population calcium response during locomotion. In a few seconds, it transitioned to phase II when the events grew and merged, ultimately forming a superevent covering the entire astrocytic network territory. The phase II corresponded to <ΔF/F> and active area peaks. Its duration correlated with the duration of locomotion. When locomotion ended, the superevent fragmented into smaller events, which then terminated. These processes determined phase III, which was followed by phase IV, termed the afterburst. The afterburst was associated with continued calcium activity in astrocytes, such as the appearance of new events. We speculate that the afterburst can be caused by the accumulation of inositol-3-phosphate (IP3) or calcium in the cytoplasm during locomotion, which can trigger calcium release from the endoplasmic reticulum. Alternatively, it could represent a response to continuing activity of adrenergic fibres that was reported to outlast the locomotion for several seconds ^28^.

Temporal integration of noradrenaline signalling by astrocytes was previously described for the different durations of optogenetic stimulation of adrenergic projections into the cortex ^27^. Oe and coauthors showed that prolonged stimulation of adrenergic fibres increases both the <ΔF/F> of population calcium response and the number of recruited astrocytes. <ΔF/F> rapidly increased for stimulation durations lasting over three seconds. The number of activated astrocytes approached 100% for stimulation durations lasting over five seconds. This result suggests that the temporal threshold of astrocytic calcium activity may be associated with the duration of noradrenaline release. In fact, a similar effect can be observed under different physiological conditions. For example, locomotion episodes are associated with continuous activation of adrenergic cortical projections ^28^. Indeed, we found that locomotion duration is also linked to the temporal threshold of astrocytic calcium response. Our results show that the animal must run for about five seconds to reach a 50% chance of suprathreshold response generation. This time is consistent with a previous report demonstrating that the integration time for astrocytic calcium response to a visual stimulus is also about five seconds ^48^.

Analysis of calcium response phases revealed the origin of the five-second temporal threshold. We found that all episodes of locomotion, regardless of their duration, triggered phase I, the emergence of new calcium events, which was identified by the IS peak. However, the superevent phase II did not develop in the case of short locomotion episodes. Since this phase largely determines the population parameters, <ΔF/F> and active area appeared very small. The animal had to overrun the IS peak for calcium events to start merging into a superevent. Five seconds is approximately the time when phase I ends, and phase II starts.

At the level of individual astrocytes, the temporal threshold of population response may be linked to the temporal threshold of somatic calcium surge ^11–13^. Somatic calcium surge is a large calcium transient in astrocytic soma that results from integration of calcium activity starting at the periphery of astrocytic anatomical domain at the beginning of locomotion. In our previous study, we reported a 4.7-second median latency between the appearance of calcium on the periphery of the astrocyte and the somatic surge ^13^. Rupprecht and colleagues found that calcium events in most distal processes occur approximately 6.5 seconds before the somatic surge ^12^. Lines and colleagues described a spatial threshold of somatic surge ^11^. They demonstrated that somatic surge requires activation of 23% of astrocytic branches, which also develops gradually over time. Thus, the integration time of calcium activity in single astrocytes is indeed comparable to the temporal threshold of the population response.

Apart from the distinction in the superevent phase II, the other phases were well pronounced in both subthreshold and suprathreshold responses. However, statistically, fewer calcium events emerged in phase I in subthreshold responses. A parsimonious explanation is that the locomotion episode was too short for this phase to develop in full. Somewhat unexpectedly, we did not observe a significant difference in the phase of event fragmentation and afterburst. This result suggests that these phases are independent of the locomotion duration and may be regulated by other mechanisms. We speculate that the afterburst may have a different physiological meaning in complex calcium dynamics in astrocytes. The superevent is a global response, a phase of singularity. During the superevent, the system cannot be described in terms of individual calcium events. During an afterburst, to the contrary, we observed the appearance of spatially distributed calcium events that could be involved in the regulation of local processes such as synaptic plasticity ^39^.

Although the superevent phase II cannot be described by the distribution of calcium events, it is characterised by a specific pattern of ΔF/F_max_ reproducible from one episode of locomotion to another. Typically, calcium elevations started in the same hotspots with the highest ΔF/F_max_ during the superevent. These hotspots also had the highest rate of calcium event initiation during quiescent periods. Mechanistically, hotspots may be linked to astrocyte morphological properties such as the surface-to-volume ratio of astrocytic processes ^53,54^, the subcellular distribution of organelles (e.g., endoplasmic reticulum and mitochondria) ^55–57^, proximity and efficacy of synapses (local and subcortical projections) ^58^ (for review see ^1^). The functional role of calcium hotspots in the astrocytic network requires further investigation. We speculate that the pattern formed by the hotspots may serve as the foundation of the astrocytic part of memory engrams through patterned cFos expression ^2,3^.

The reproducibility of the population calcium response pattern from one locomotion to another suggests that the response may be described as an evolution of a low-dimensional deterministic dynamical system. Indeed, the results of DMDc modelling confirmed this hypothesis. The model, fitted to pairs of directly consecutive ΔF/F frames and instantaneous speed and acceleration values, predicts the characteristic phases in the response evolution, temporal threshold, pattern of hotspots, etc, thus providing insight into intrinsic properties of the biophysical mechanisms of calcium dynamics. The A (dependence on previous state) and B (dependence on control) operators, which were fitted in the model, are responsible for alterations of the response and hence can represent biologically relevant factors such as animal state (e.g. vigilance, stress, prior activity) and the state of BAM (e.g., plasticity of astrocytes, neurons, subcortical projections).

Despite the striking similarity between the calcium response in a real astrocytic network and the response simulated with DMDc, we observed several differences that may be attributed to the nonlinearity of the calcium response. The <ΔF/F> elevation in the DMDc model did not show a decline, while the afterburst phase had some oscillatory modes. In experimental data, <ΔF/F> increased at the beginning of the superevent phase II and slowly declined towards the end of the locomotion episode. Indeed, the decline and the refractoriness of astrocytic calcium response during locomotion have been previously reported ^13,16^. This phenomenon may play a role in preventing calcium excitotoxicity.

In summary, this study characterises the population calcium response in cortical astrocytes during locomotion as a reproducible, multiphase process initiated by the rapid emergence of new calcium events. We demonstrate that a full-scale network response, involving the growth and merging of these events into a superevent, requires locomotion to persist beyond a critical five-second temporal threshold. This timing likely reflects the integration period needed to trigger somatic calcium surges within individual astrocytes. Spatially, the superevent pattern originates from consistent and reproducible hotspots, which also serve as activity hubs in quiescent periods. Our DMDc modelling successfully captures these complex spatiotemporal dynamics, revealing that the response can be described as a low-dimensional deterministic system whose emergent properties, including the characteristic phases and temporal threshold, are shaped by biological factors like brain state and plasticity ^59^.

## Methods

### Animals

All procedures were performed in accordance with local guidelines for animal care, following protocols approved by both the Institute of Bioorganic Chemistry’s ethical committee and the animal ethics and welfare committee of Jiaxing University. These protocols are based on FELASA guidelines and recommendations. For the 2-photon imaging experiments, we used male C57BL/6 mice aged 4 to 8 months. The mice were housed in groups under standard conditions, maintained on a reversed 12-hour light/12-hour dark cycle (with lights off at 9 AM), and had *ad libitum* access to food and water.

### Surgical procedures

The surgeries were conducted on mice aged 3 to 4 months. During the surgical procedures, the mice were anesthetised with isoflurane (3% in O_2_ for induction, 0.8 – 1.5% for maintenance) while being placed in a stereotaxic frame (RWD, China). The body temperature of the animals was maintained at 37 °C using a heating pad. An eye ointment (Vidisic, Germany) was applied to keep the eyes moist throughout the surgery. The head skin was removed, and a 3-mm craniotomy was performed above the right primary somatosensory cortex (rS1) using a dental drill (RWD, China). During the surgery, the skull was kept moist through the application of sterile saline. AAV9-gfaABC1D-cyto-GCaMP6f (1 µl, titre 10^11^ vg/mL) was delivered via stereotaxic injection using a 5 µL glass pipette (Drummond, USA) to rS1 (AP −2.3, ML +0.5 from bregma, and DV −0.8, −0.6, −0.4, −0.2 from the surface of the dura) to achieve astrocyte-specific expression of the genetically encoded calcium indicator GCaMP6f. The plasmid pZac2.1 gfaABC1D-cyto-GCaMP6f for virus production was a gift from Baljit Khakh (Addgene plasmid # 52925; http://n2t.net/addgene:52925; RRID: Addgene_52925) ^60^. After each injection, the pipette was left in place for at least 5 minutes to ensure proper diffusion of the AAV and prevent reflux. Following the AAV injection, a 5-mm round cover glass (Thomas Scientific, USA) and a lightweight, custom-made stainless-steel head plate were affixed to the skull over the craniotomy using a mixture of dental cement (Simplex Rapid Powder, Kemdent, UK) and cyanoacrylate glue (Cosmofen Ca-12, Germany) for head fixation in the experimental setup. An analgesic (ketoprofen, 2.5 mg/kg) was administered subcutaneously to the mice after surgery.

### Behavioural training and two-photon imaging

Experiments commenced after a recovery period of 4 to 6 weeks, along with 1 week for habituation to the experimental setup and imaging sessions. Habituation included animal handling for 10 to 15 minutes daily over 5 days. Subsequently, the mice were acclimated to being head-fixed under the objective of a multiphoton fluorescence microscope (Femtonics, FemtoSmart, Hungary) and running on a freely rotating disk (Gramophone, Femtonics). All experiments were conducted between 10 AM and 6 PM. Mice were imaged using an Olympus 20×1.0 NA water-immersion objective (model N20X-PFH 20X, Olympus). A Ti:sapphire laser (Chameleon Ultra, USA) emitting at 920 nm was used to excite the fluorescence of GCaMP6f. The GCaMP6f fluorescence was isolated using a 520/60 nm bandpass filter (green channel) while the autofluorescence was isolated using a 605/70 nm bandpass filter (red channel). The signals from both channels were detected with GaAsP photomultipliers H10770PA-40 (Hamamatsu, Japan). Images (512 × 512 pixels, 550 × 550 μm) were acquired using a resonant scanner at a rate of 31 frames per second (fps) in layer 1 of rS1. Simultaneously, the rotation of the Gramophone disk was monitored using locomotion-tracking software to assess the animals’ speed. All sessions were recorded in infrared light using an ELP 1.3-megapixel USB camera (Shenzhen Ailipu Technology Co., Ltd, China) at 20 fps. Each experimental session did not exceed 2 hours. The Gramophone disk was cleaned with 70% ethanol and distilled water between sessions.

### Data analysis

To enhance the signal-to-noise ratio, raw frames captured at 31 fps were summed over consecutive, non-overlapping batches of 10 frames (10x temporal binning), resulting in an effective frame rate of 3.1 fps. After binning, the pipeline for data analysis included: correction for lateral shifts (motion correction), denoising, calculation of a slowly varying baseline for each pixel, ΔF/F calculation for each pixel, application of ΔF/F thresholds to identify active pixels, and analysis of suprathreshold regions in each frame.

### Motion correction

The correction of lateral translational motion between frames was done with registration.phase_cross_correlation function from the [scikit-image] Python library ^61^. Changes in GCaMP6f could affect image correction algorithms in the green channel. Therefore, the autofluorescence signal recorded in the red channel was used to estimate lateral shifts between frames (Extended Data Figure 7). Then the obtained shifts were applied to correct motion in the green channel. We applied the following preprocessing for the binned red channel frames: 1. Each frame was represented as a linear combination of 50 principal frames by the truncated singular value decomposition (tSVD) on flattened frames. The principal frames were spatially smoothed with a Gaussian filter (σ = 1 px), while the time series of coefficients for each of the principal frames were temporally smoothed by applying a 3-point median filter followed by a Gaussian filter (σ = 0.5 frames), after which the inverse SVD and reshaping produced final filtered frames. Empirically, this transform resulted in spatially smoothed denoised frames with dampened temporal variations, facilitating the calculation of spatial shifts. To provide a template to align all frames, the frames were clustered using the self-organising maps algorithm (grid size, 1 × N_f_ / 100, where N_f_ is the number of frames), and the centre of the cluster with the most elements was used as the template. The obtained lateral shifts for each frame were then applied to the noisy binned frames of the green channel.

### Denoising

To further increase the signal-to-noise ratio of the images, we used an in-house SVD-based denoising algorithm. The raw recording of the size N_f_ × 512 × 512, where N_f_ is the number of frames, was divided into 8 × 8 px spatial blocks with 4 px overlap. The data within each block were flattened to produce N_f_ × 64 matrices, which were then factorised with truncated SVD, keeping only a few first components. Each spatial component was reshaped back to an 8 × 8 matrix and processed with a 3 × 3 median filter. The motivation for doing SVD in small blocks rather than full frames was to avoid missing local small-amplitude dynamics, which would not contribute to global variance in the whole frame. At this first stage, temporal SVD components were still noisy, but we reasoned that if neighbouring blocks had similar dynamics, they would share similar parts in the temporal components. On the second stage, temporal components within a 100 × 100 px neighbourhood were collected and clustered within a sliding 50-frame temporal window using agglomerative clustering with Ward’s linkage. Next, within each window, the filtered estimates of the temporal components were taken as cluster centres. Estimates in window overlaps were averaged. Finally, an inverse SVD decomposition using filtered spatial and temporal components provided estimates of the signal within each block; estimates in the overlapping parts of the blocks were averaged.

Due to the low photon count and stochastic nature of photon emission, the noise of the fluorescent signal is characterised by a shifted and stretched Poisson distribution. Because our denoising scheme assumes Gaussian noise, we applied a variance-stabilising Anscombe transform before the denoising stage and an optimal inverse Anscombe transform after denoising ^62^.

### Calculation of the slowly varying baseline and of ΔF/F

The slowly varying baseline was calculated in each pixel. Firstly, the signal was filtered by a median filter with a 3-second window, followed by Gaussian smoothing with σ = 1.5 s. Then, a linear interpolation was drawn through those local minima in the filtered signal, which lie within 1.5 standard deviations of signal noise (estimated as the standard deviation of the first derivative of the signal divided by √2) from the greyscale morphological closing of the greyscale morphological opening of the filtered signal. Next, the baseline estimate was Gaussian-smoothed with σ = 15 s. Finally, to compensate for bias resulting from fitting to local minima, the baseline was shifted to bring the mode of the residuals from the original signal towards zero. The resulting baseline (*F_0_*) was calculated in each pixel independently and was used to define ΔF/F:

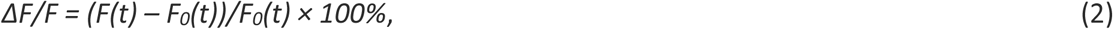

where *F(t)* is the value of fluorescence at a specific point in time, and F0(t) is the baseline calculated for this point in time.

Because GCaMP6f was expressed only in GFAP-expressing astrocytes, some areas within the field of view, occupied by the other cell types, remained unstained and were excluded from the analysis. Unstained areas were defined in a time-averaged baseline frame <F_0_(t)>. We estimated the first mode *M* of the pixel <F_0_(t)> distribution. Then the pixels with <F_0_(t)> < (*M* – SD) were labelled as unstained. <ΔF/F>, active area and segment density were calculated within the stained areas.

### Calculation of activity segments and segment-based parameters

A ΔF/F threshold of 15% was chosen empirically because it yielded the most visually informative representations of activity. Suprathreshold pixel clusters larger than 16 pixels within a frame were defined as segments if they overlapped with corresponding clusters in at least three consecutive frames. For each frame, the fractional area occupied by active segments, normalised to the total stained area within the field of view, was used as a population parameter of calcium activity and referred to as the active area.

Segments with no overlap with the preceding frame were classified as initial segments (IS), those with no overlap with the subsequent frame as terminal segments (TS), and all remaining segments as middle segments (MS). Segment density was calculated as the number of segments per total stained area. We also quantified segment size distributions for each segment type and used the mean segment size, pooled across types, as another indicator of calcium activity.

### Detection of locomotion episodes

A speed threshold of 0.5 cm/s was used to detect the onset of a locomotion episode. During an episode of locomotion, the animal ran at a variable speed. If the speed decreased below the threshold (but not below 0.25 cm/s) for less than 3 s, this interval was included in the locomotion episode. Short animal moves (less than 3 s) were not considered as episodes of locomotion.

### Data-driven modelling of astrocytic calcium responses

We used the pydmd (https://github.com/PyDMD/PyDMD) module to fit dynamic mode decomposition with control model (DMDc) to experimental data (Extended Data Figure 8) ^63,64^. The estimates for the *A* and *B* matrices, together with initial conditions *F(t = 0)*, allowed for the simulation of system dynamics in response to a real or synthetic sequence of control parameters. Notably, for high-dimensional systems, matrix A can be prohibitively large, and for practical reasons, it was approximated by a set of its eigenvectors.

Consecutive pairs of ΔF/F frames were used as state snapshots, while instantaneous mouse speed and acceleration (numerical first derivative of speed) served as control parameters (Extended Data Figure 5). However, using full ΔF/F frames was impractical computationally and numerically unstable. Instead, we applied DMDc to the first 50 temporal components of the SVD decomposition of ΔF/F data. Simulation results for synthetic or real mouse speed profiles were then transformed back into ΔF/F space by inverse SVD transform. Applying DMDc to SVD components as opposed to applying it directly to ΔF/F data did not reveal any noticeable differences in the solution.

DMD algorithm is sensitive to noise and may predict spurious unstable oscillatory modes. Response characteristics in the resulting model depended on the data samples used for fitting and, to a lesser extent, on initial conditions. In some cases, the model dynamics can become unstable even after the speed has returned to zero. To reduce the influence of noise, an ensemble approach to DMD, similar to the idea of bagging DMD was used ^65^: for a given imaging session data, a batch of 50 DMDc models was fitted on randomly selected subsets of temporal point pairs, after which the responses of each model in the batch were averaged to obtain the final model prediction.

In simulations, we used both real locomotion episodes and synthetic speed profiles as control parameters. Synthetic speed profiles were parametrised with three parameters: maximal speed, maximal acceleration, and duration, and were modelled as a product of two hyperbolic tangent functions:

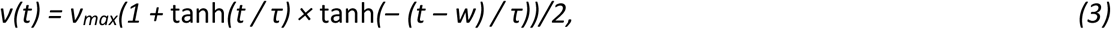

where *v_max_* is maximal speed, *w* is locomotion duration, and *τ = (v_max_/a_max)_)/2* describes the steepness of the rise and fall of speed such that peak acceleration is equal to *a_max_ (*Extended Data Figure 5). Based on the available dataset of experimental locomotion episodes, we fixed *v_max_= 4.0 cm/s, a_max_ = 3.2 cm/s^2^* as the reference values of the corresponding variables (median, [iqr] for speed: 4.0 [2.3, 5.6] cm/s, for acceleration: 3.2 [1.8, 5.9] cm/s^2^) and varied the duration of the synthetic locomotion episodes.

### Statistical Analysis

Due to the nested nature of the data (i.e., multiple locomotion episodes per imaging session and multiple imaging sessions per mouse), we employed linear mixed-effects models (LMM) as the primary statistical tool. Specific animal and imaging session numbers were treated as random effects, while comparison groups were treated as fixed effects. Statsmodels Python module was used to fit statistical models ^66^. Data presented as M [Q1; Q3] ± 1.5 IQR, p-values are LMM; n-numbers were defined and given in the figure legends. The experimenters were not blinded to the experimental conditions, and no randomisation was performed.

---

**Extended Data Figure 1:**
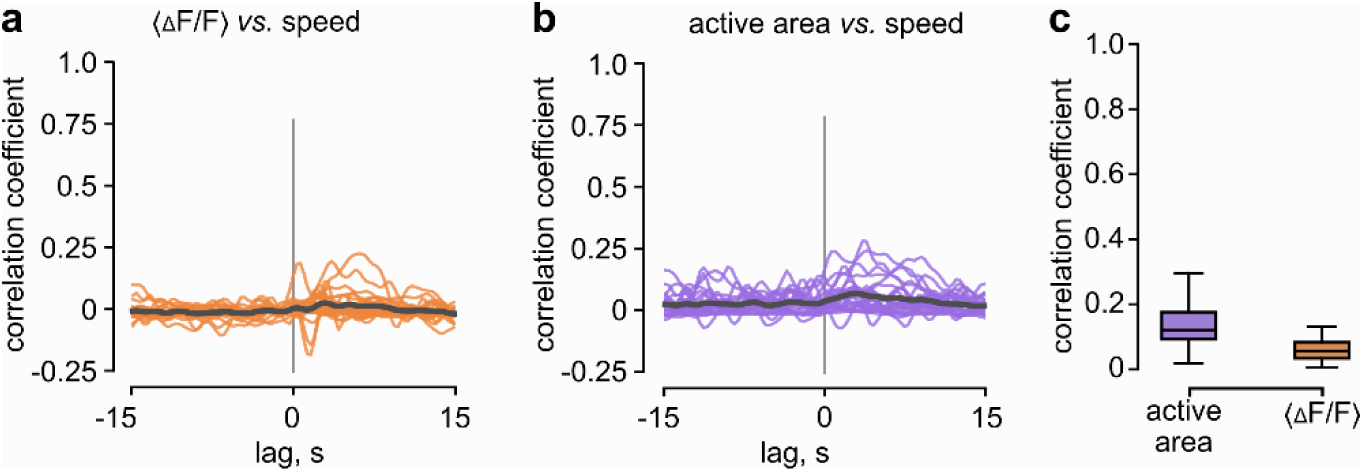
Calcium activity fluctuations during mouse quiescence have little correlation with short movements. a, Cross-correlation graphs between <ΔF/F> and mouse speed during quiescence. **b,** Cross-correlation graphs between active area and mouse speed during quiescence. **c,** Summary of correlation coefficients for cross-correlations at panels *a* and *b*. Data presented as median [Q1; Q3] ± 1.5 IQR, *n* = 26 periods of quiescence, 12 imaging sessions from 3 animals.

**Extended Data Figure 2:**
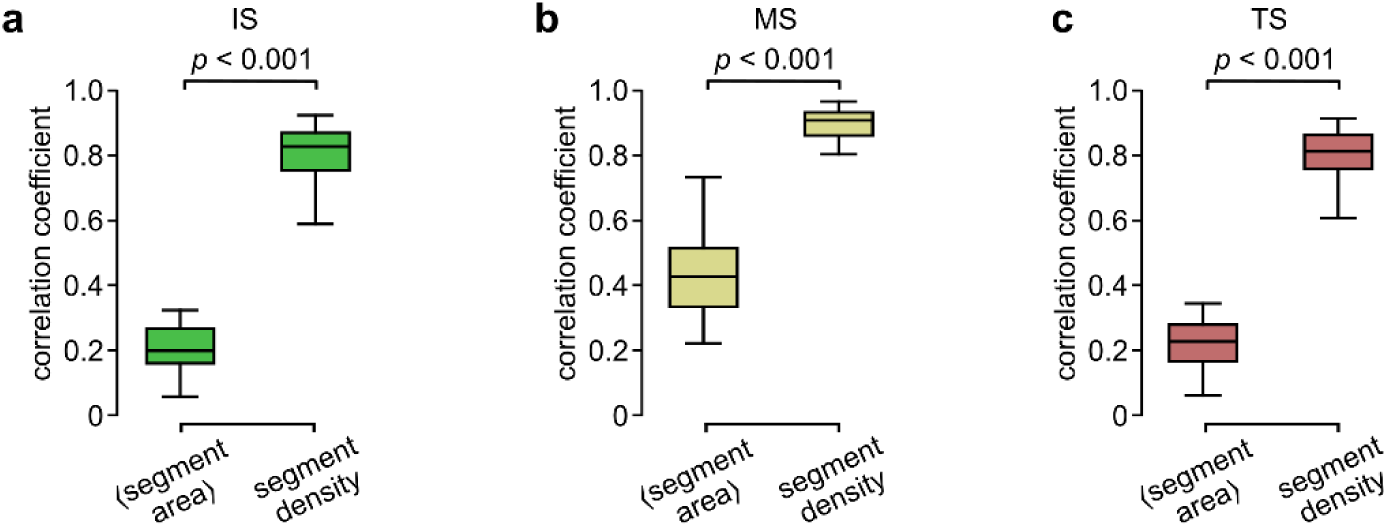
Calcium activity fluctuations during mouse quiescence have a higher correlation with segment density than <segment area>. Summary data of correlation coefficients between <segment area> and active area, and correlation coefficients between segment density and active area for IS (a), MS (b), and TS (c). Data presented as median [Q1; Q3] ± 1.5 IQR, *p*-values are Mann-Whitney U test, *n* = 26 periods of quiescence, 12 imaging sessions from 3 animals.

**Extended Data Figure 3:**
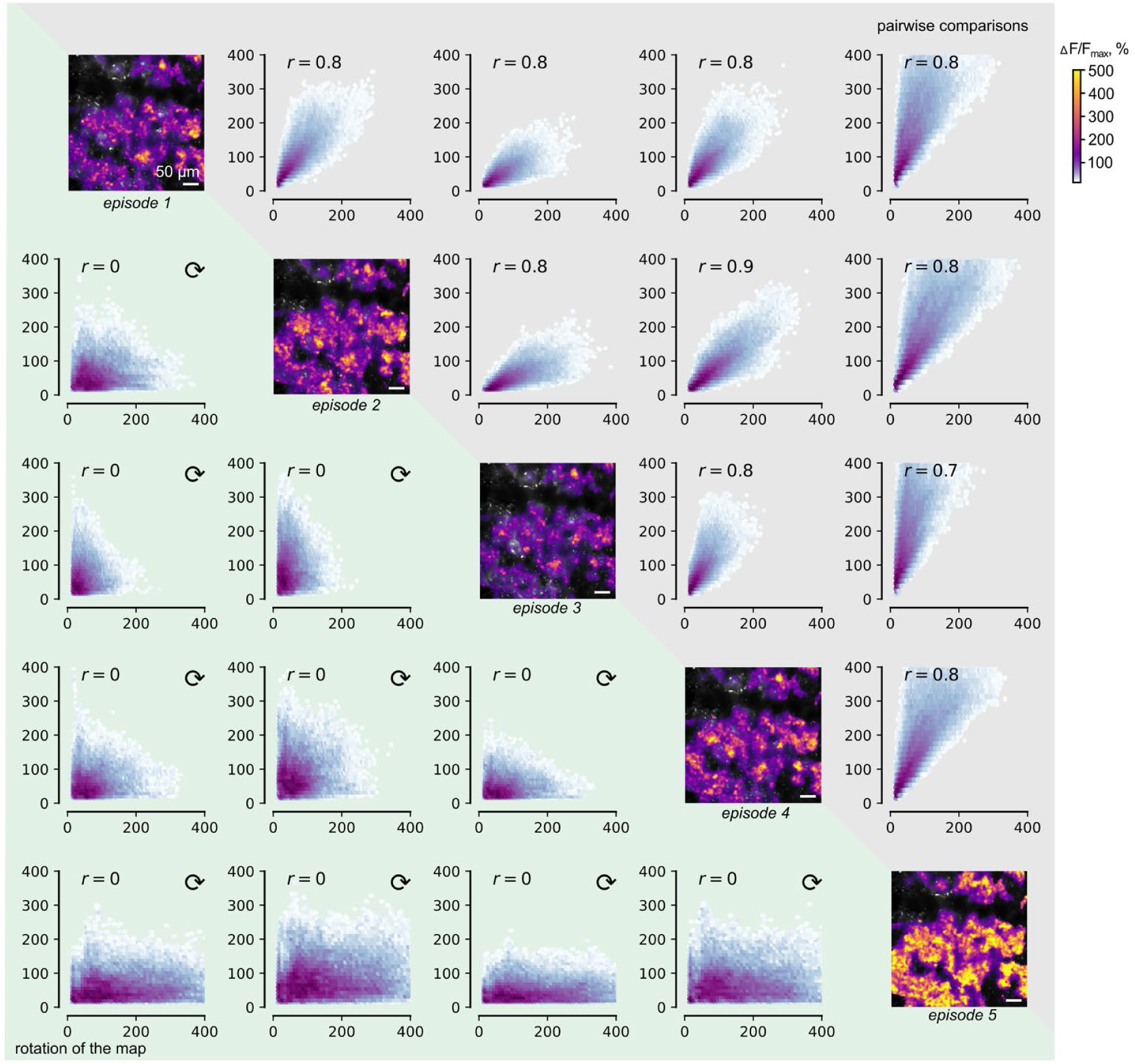
Maps of ΔF/Fmax of calcium transients in five sequential episodes of locomotion are shown along the diagonal. The relationships between ΔF/Fmax in corresponding pixels between pairs of episodes of locomotion are shown in the upper right corner. The same relationships, when one map in each pair was rotated by 90 degrees (random correlations), are shown in the lower left corner.

**Extended Data Figure 4:**
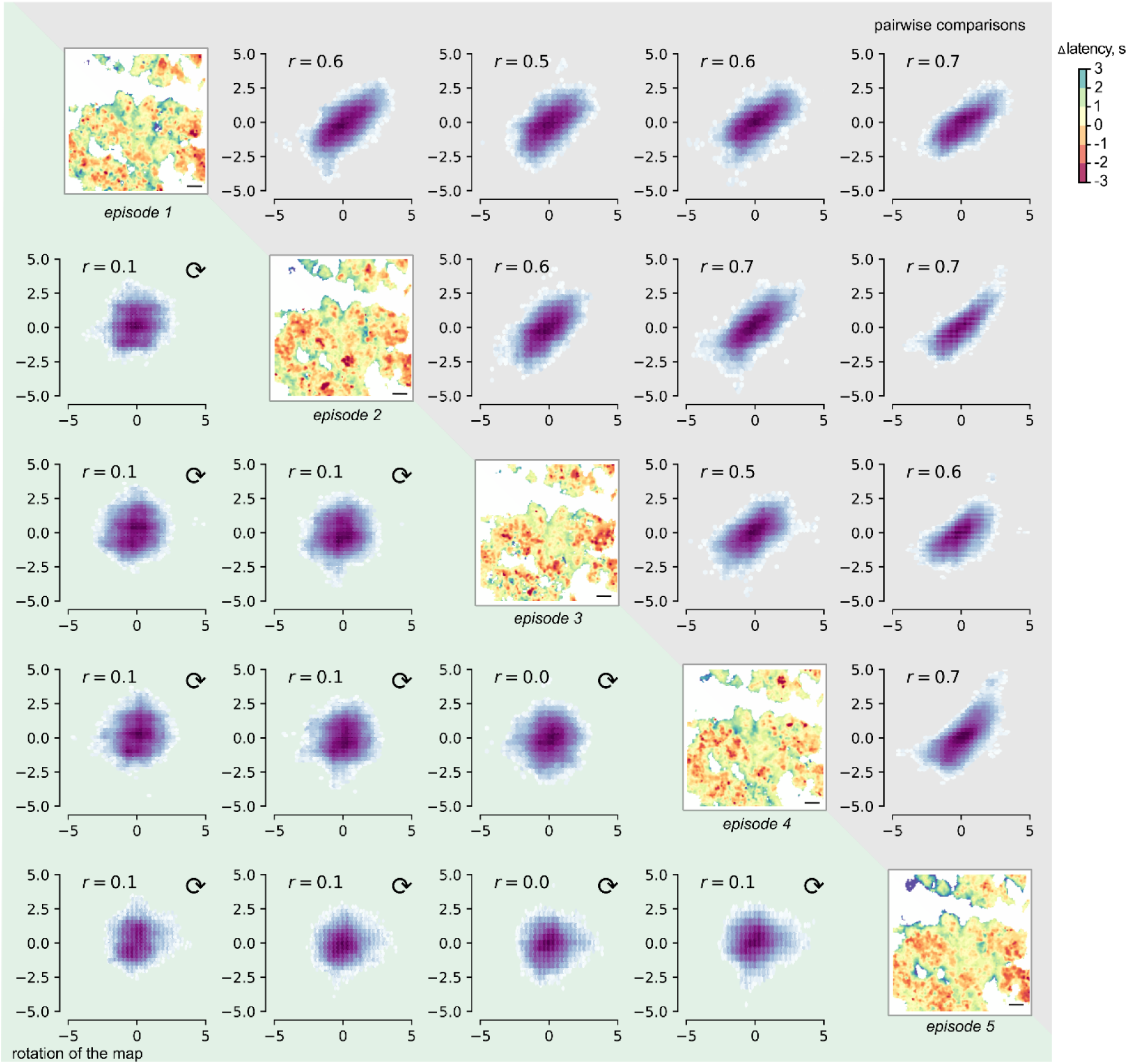
Correlations between pairs of Δ latency maps for different locomotion episodes in the same imaging session. Maps of Δ latencies of calcium transients in five sequential episodes of locomotion are shown along the diagonal. The relationships between Δ latencies in corresponding pixels between pairs of episodes of locomotion are shown in the upper right corner. The same relationships, when one map in each pair was rotated by 90 degrees (random correlations), are shown in the lower left corner.

**Extended Data Figure 5:**
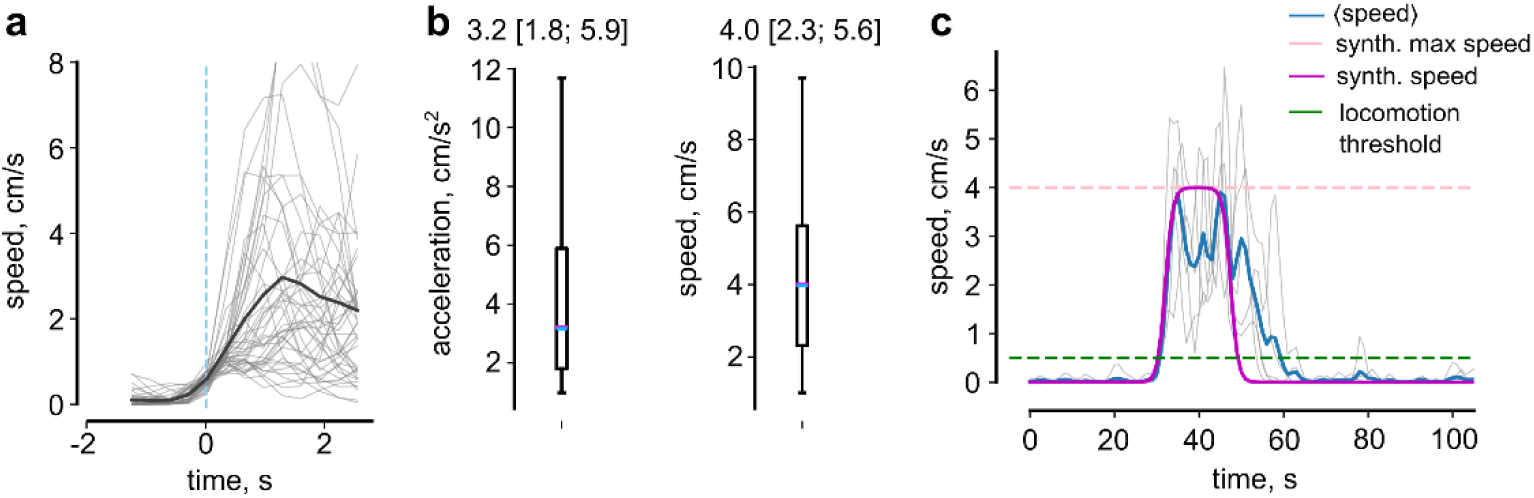
Calculation of control parameter C(*t*) for DMDc model. **a.** Multiple episodes of locomotion aligned by their starting points (blue dashed line). Thin grey lines indicate the mouse speed during multiple locomotion episodes. Thick black line – mean locomotion speed. **b.** The summary of mouse acceleration (*left*) and speed (*right*). **c.** Multiple locomotion episodes (grey), mean curve (blue) and synthetic speed for DMDc model (violet).

**Extended Data Figure 6:**
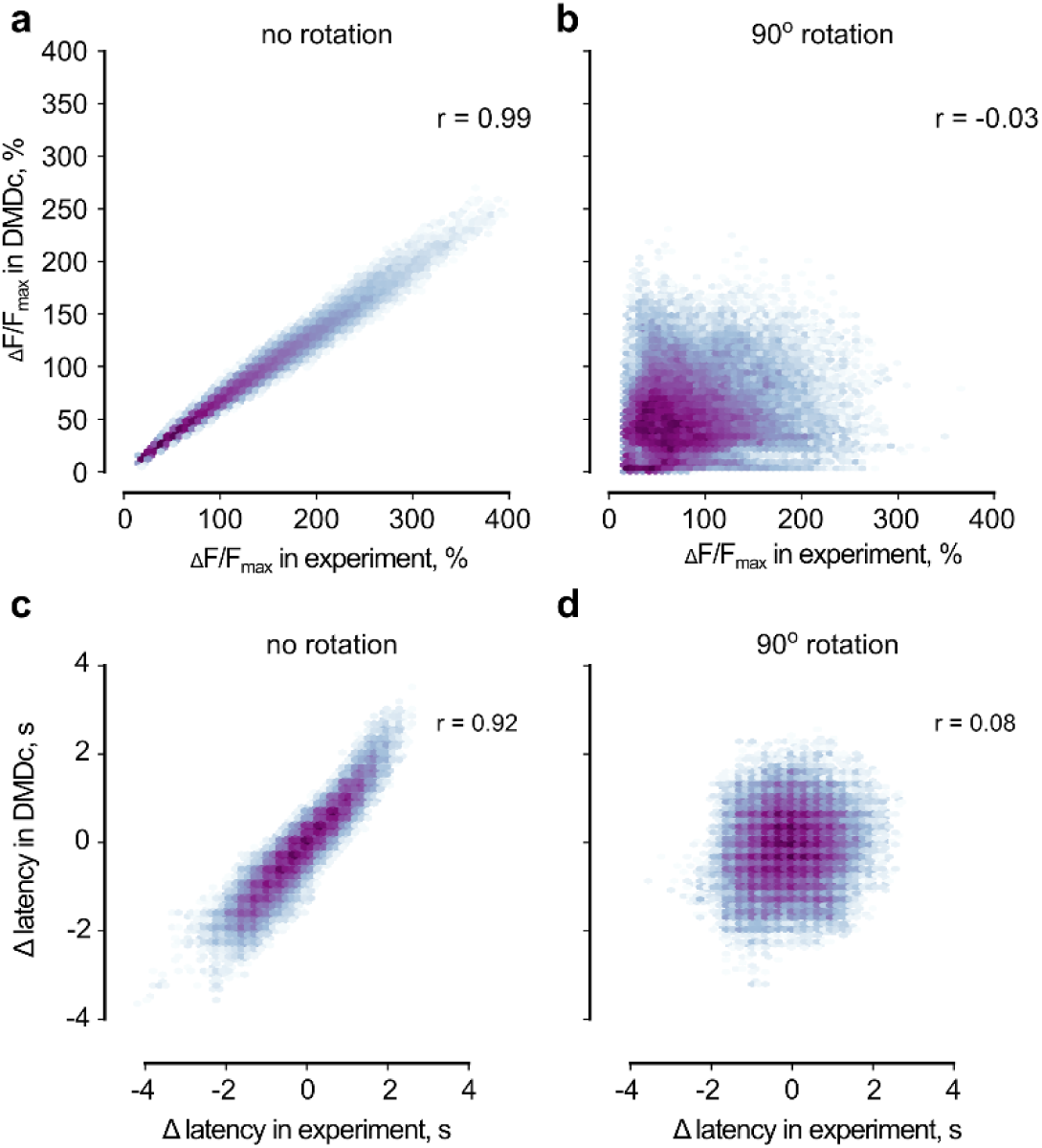
Correlations between pairs of ΔF/Fmax maps and Δ latency maps for experimental and synthetic data. **a.** The relationship between ΔF/Fmax in corresponding pixels in the experiment and in the DMDc model in response to the same locomotion profile. ‘r’ is Pearson’s correlation coefficient. **b.** Same relationship as in panel *a*, when one map was rotated by 90 degrees. **c.** The relationship between Δ latency in corresponding pixels in the experiment and in the DMDc model in response to the same locomotion profile. ‘r’ is Pearson’s correlation coefficient. **d.** Same relationship as in panel *c*, when one map was rotated by 90 degrees.

**Extended Data Figure 7:**
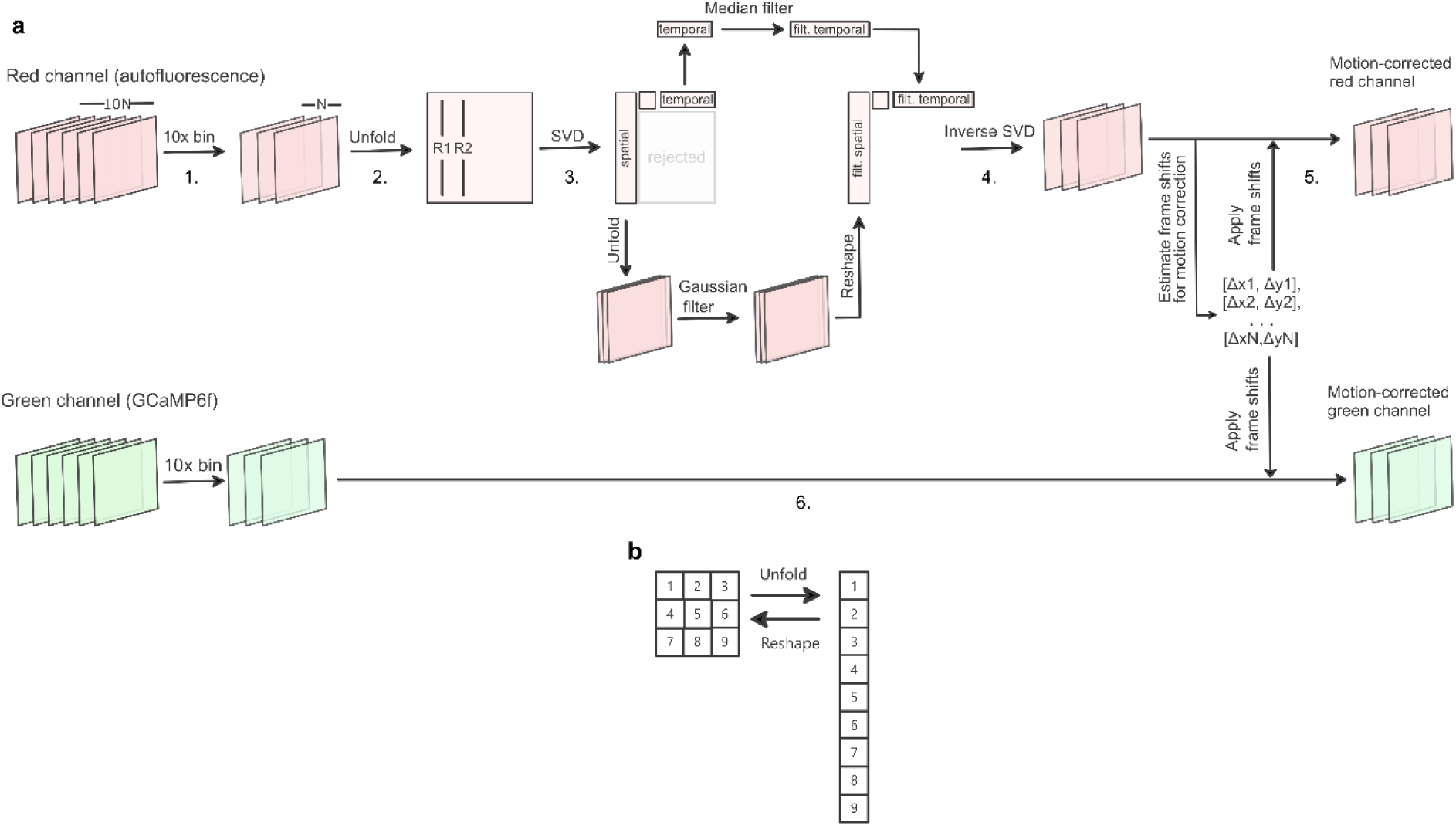
Image preprocessing: motion correction pipeline. **a.** The goal is to partially denoise and smooth red-channel frames, calculate frame shifts that would correct frame displacements in the red channel and apply them to the corresponding frames in the green channel. Below are the detailed steps: Red channel: 1. Raw frames are 10× binned. 2. Each frame in the binned record is unfolded to a vector. If the frame was of the size R × C, then the total number of pixels P = RC (rows times columns), resulting in a P × N matrix. 3. The matrix is then decomposed using truncated singular vector decomposition, using Numpy slicing notation: *P ≈ U[:,:k]* Σ*[:k,:k] V’[:k,:]*, where first *k* left (*U*) and right (*V’)* singular vectors and first *k* elements of the diagonal matrix Σ are kept, and the rest are discarded can be interpreted as unfolded principal frames; the columns of U can be interpreted as unfolded principal frames, and, after reshaping them to R × C images, they are Gaussian-blurred and after unfolding are next used in place of original columns of *U (3’);* the rows of *V’* can be interpreted as principal temporal signals, so they are filtered with a median filter to get rid of sharp peaks and used in place of original rows of *V’ (3’’)* 4. Inverse SVD with the filtered columns of U and filtered rows of V*’* gives an adaptively denoised version of the record in the red channel. 5. All denoised red-channel frames are aligned to correct for motion shifts, and pixel position transforms for each frame are recorded. 6. The obtained pixel position transforms are applied to 10× binned green channel frames to produce a motion-corrected green-channel recording. **b.** Illustration of the unfolding of the frame to a vector on step 2 of the preprocessing pipeline.

**Extended Data Figure 8:**
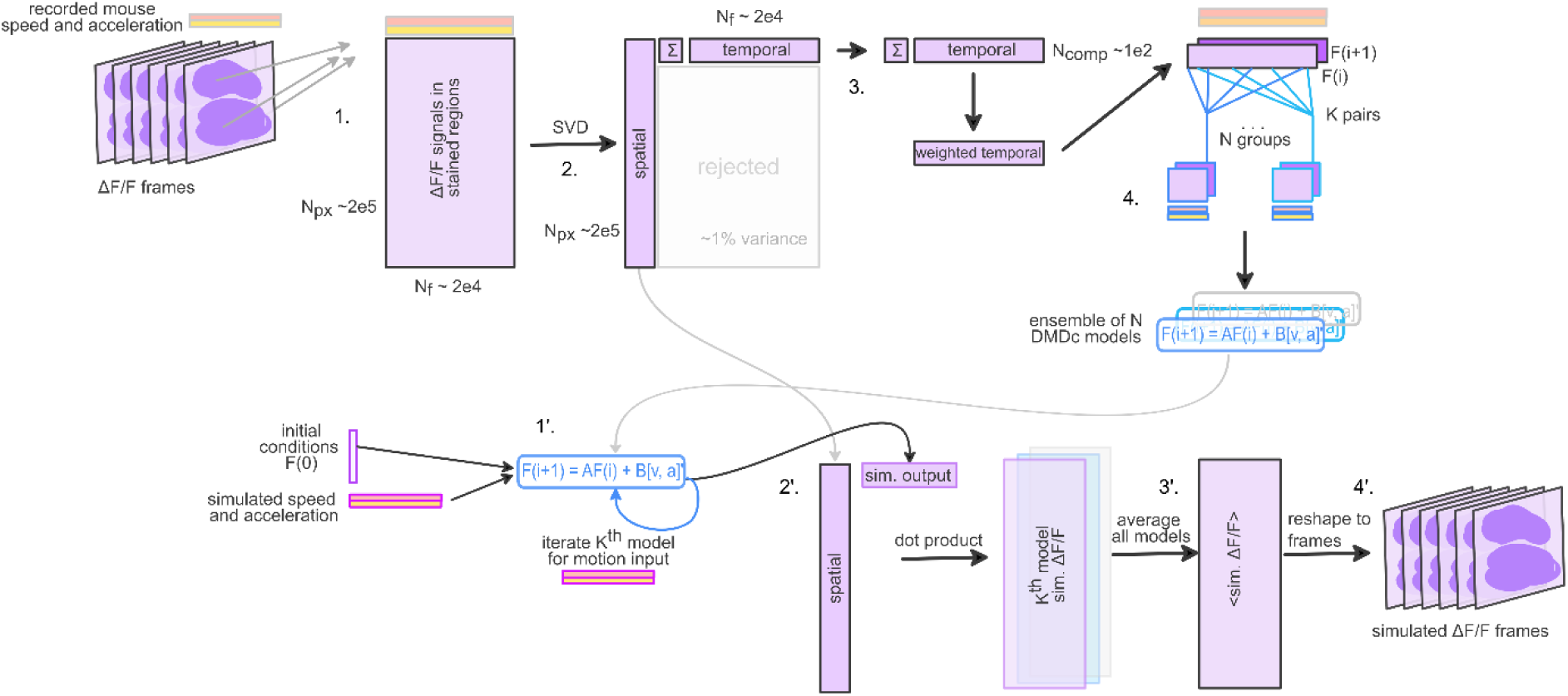
DMDc application to fluorescence dynamics. The goal is to fit a DMDc model to the dependence of experimental ΔF/F data on animal speed and numerically computed acceleration. The DMDc model can then predict ΔF/F responses from an initial ΔF/F value and speed + acceleration profile. *Upper row:* fitting an ensemble of DMDc models. 1. In each frame, ΔF/F values from all stained pixels are unfolded to a vector, resulting in a tall and skinny matrix *P* of size Npx × Nf. 2. Truncated singular value decomposition (SVD) approximates this matrix with a dot product of k spatial and k temporal components, or using Numpy slicing notation: *P ≈ U[:,:k]*Σ*[:k,:k]V’[:k,:],* where first k columns of U can be interpreted as unfolded principal ΔF/F frames, and first k rows of V*’* can be interpreted as the principal time signals, which we would like to fit with the model. The parameter k is typically 50…100 and set to capture around 95…99% of the variance in the original matrix P. 3. Note that U and V* are unitary, thus their corresponding columns and rows are normalised to 1, while all magnitude information is stored in the diagonal matrix Σ. To weigh the importance of each principal time signal, we further work with the *ΣV\** product. 4. The DMDc model fits matrices *A* and *B* to be able to predict the state of the dynamical system *F* at time point *j + 1* given the state at time point *j* and the value of the control parameter (the pair of speed and acceleration values) at time *j*: *F(j + 1) ≈ AF(j) + B[v(j), a(j)]’* for any time moment *j*. To mitigate the effect of noise, we randomly chose K groups of N (j,j+1) time point pairs and independently fitted an ensemble of K DMDc models. *Lower row:* simulation with DMDc models. 1’. Starting with initial conditions F(0) and simulated speed + acceleration profile, iteratively applying the K DMDc models from the ensemble to obtain simulated weighted temporal components. 2’. Using dot-product with the spatial components to obtain predictions of ΔF/F signals and 3’. averaging them over the ensemble. 4’. Reshaping the frames to obtain the simulated ΔF/F dynamics for the given locomotion profile.

